# C_4_ Grasses Employ Distinct Strategies to Acclimate Rubisco Activase to Heat Stress

**DOI:** 10.1101/2022.08.24.505184

**Authors:** Sarah C Stainbrook, Lindsey N Aubuchon, Amanda Chen, Emily Johnson, Audrey Si, Laila Walton, Angela Ahrendt, Daniela Strenkert, Joseph Jez

**Author notes:** **Correspondence:** Sarah Stainbrook.

## Abstract

Rising temperatures due to the current climate crisis will have devastating impacts on crop performance and resilience in the near future. One key step that limits plant photosynthetic performance under higher temperatures is the activity of the thermolabile enzyme rubisco activase (RCA). RCA is highly conserved in photosynthetic organisms, including C_4_ crops such as *Zea mays* (maize) and *Sorghum bicolor* (sorghum) which are crucial components of global food supply and the bioenergy sector. While rubisco is the most abundant protein on earth and responsible for carbon fixation, RCA is an essential chaperone required to remove inhibitory sugar phosphates from the active site of rubisco to allow for continued CO_2_ fixation. We set out to understand temperature-dependent RCA regulation in four different C_4_ plants, with a focus on the crop plants maize (two cultivars) and sorghum, as well as the model grass *Setaria viridis* (setaria). Gas exchange measurements confirm that CO_2_ assimilation is indeed limited by Ribulose 1,5-bisphosphate (RuBP) carboxylation in these organisms and at high temperatures. All three species express distinct sets of RCA isoforms and each species alters the isoform and proteoform abundances in response to heat; however, the changes are species-specific. In order to understand how even subtle changes in the molecular environment of the chloroplast stroma affect RCA function during heat acclimation, we examined the regulation of RCA activity directly with respect thermostability, the ratio of ADP to ATP and the concentration of Mg^2+^ ions. As shown previously, the activity of RCA is modulated by a combination of these variables, but surprisingly, how these biochemical environment factors affect RCA function differs vastly between the different C_4_ species, and differences are even apparent between different cultivars within a single species, both with respect to proteoform abundance and regulation. Our results suggest that each grass evolved different parts of the RCA regulation portfolio and we conclude that a successful engineering approach aimed at improving carbon capture in C4 grasses will need to accommodate these individual regulatory mechanisms.

## 6 Introduction

Ribulose-1,5-bisphosphate carboxylase/oxygenase (rubisco) is responsible for almost all carbon fixation on Earth. Rubisco catalyzes the addition of CO_2_ to the C5 sugar ribulose-1,5-bisphosphate (RuBP), producing two molecules of glycerate-3-phosphate, which are subsequently further assimilated in the Clavin-Benson-Bassham (CBB) cycle. Rubisco is a poor enzyme; it is slow, assimilating only ∼ 3-10 CO_2_ molecules / s, lacks in specificity, both between carbon (CO_2_) and oxygen (O_2_), which induces photorespiration resulting in a further reduction of the efficiency of CO_2_ assimilation, and its C5 substate, allowing for the binding of other sugar phosphates than RuBP. Inhibitory sugar phosphates thereby directly bind to the active site of rubisco in place of the intended RuBP substrate itself, inactivating rubisco enzyme activity until the removal of the sugar phosphate. For this purpose, Rubisco activase (RCA) has evolved alongside rubisco, facilitating the removal of inhibitory substrates and thereby restoring rubisco activity, rendering this protein essential for photosynthetic carbon fixation. However, RCA itself is not without issues, especially with regard to thermotolerance; its activity is inhibited by even moderate heat (i.e. 32.5°C in maize) (Busch and Sage 2017; Steven J. Crafts-Brandner and Salvucci 2002). RCA’s inability to function properly at higher temperatures has grave implications for crop yield, especially given the rise in global average temperatures and the increase in heat waves.

Higher plants have evolved further to compensate for some of rubisco’s deficiencies. In C_3_ plants, photosynthesis takes place entirely within the mesophyll, in the vicinity of O_2_ produced by photosystem II, promoting photorespiration and its costly corrections. Photorespiration increases with rising temperatures (Sage 2013), which has led some to propose that the thermolability of RCA in C_3_ plants is a protective feature that disables photosynthesis under unfavorable conditions, similar to a fuse (Degen, Worrall, and Carmo-Silva 2020; Sharkey et al. 2001). In C_4_ plants however, rubisco functions exclusively in the bundle sheath cells, which have reduced PSII expression and activity. Rubisco is sequestered away from the primary site of PSII-dependent O_2_ production but receives CO_2_ from mesophyll cells through a pumping mechanism. This adaptation improves the efficiency of photosynthesis by limiting photorespiration to less than a fifth of that of comparable C_3_ grasses (Kanai and Edwards 1999). *In vitro*, the turnover speed of fully active rubisco improves with increasing temperature, hinting that C_4_ plants could photosynthesize more efficiently at higher temperatures if RCA could maintain rubisco activity *in vivo* (S J Crafts-Brandner and Salvucci 2000). However, the RCA expressed in C_4_ plants is still thermolabile, and inactivation of rubisco by binding of inhibitory sugar phosphates remains the predominant reason for reduced CO_2_ assimilation at elevated temperatures. The RCA of C_4_ plants is therefore an attractive target for protein engineering with the goal of improving its thermostability and thereby to increase CO_2_ assimilation when plants experience high temperatures (Parry et al. 2013).

Most plant species express two RCA isoforms, α and β, which depending on the species can arise either from alternative splicing of a single gene (e.g. spinach, Arabidopsis) or from two individual, dedicated genes (e.g. maize, setaria, sorghum). The β isoform is approximately 43 kDa in size and can undergo post-translational proteolytic cleavage to form a 41 kDa proteoform in maize (Z. Yin et al. 2014; Vargas-Suárez et al. 2004). In most species, the β isoform is expressed constitutively. The α isoform is slightly bigger, approximately 45kDa, and is mostly identical to the β isoform with the addition of a C-terminal domain (CTD). The CTD contains two cysteines capable of forming a disulfide bond which regulates RCA function according to the redox state of the chloroplast, likely by changing its affinity for ATP (D. Wang and Portis 2006; Scafaro, De Vleesschauwer, et al. 2019). Redox state dependent regulation of the α isoform couples the activity of RCA to illumination (N Zhang and Portis 1999). The α isoform of Arabidopsis has also been reported to undergo inhibitory phosphorylation at threonine 78 near the N-terminus (Kim et al. 2019). This threonine is present in the α isoforms of maize and sorghum but absent in setaria. Additional post-translational modifications have been observed but their influence on protein activity is not fully understood (Harvey et al 2022).

The α and β monomers also can differ in thermostability, with the α isoform frequently being reported as more thermostable (Steven J Crafts-Brandner et al. 1997). The thermolability of RCA may be due to structural changes such as denaturation and aggregation (Barta et al. 2010). However, RCA activity is also affected by other factors inside the chloroplast. The ATP/ADP ratio is a measure of the status of photosynthetic electron transport in the chloroplast and can affect RCA function in some species (Carmo-Silva and Salvucci 2013; Barta et al. 2010; Kallis, Ewy, and Portis 2000). In the α isoform, this sensitivity appears to be modulated by negatively charged residues in the CTD (Wang and Portis 2006), but sensitivity to elevated ADP is also observed in tobacco which lacks the α isoform and therefore the CTD (N Zhang and Portis 1999; Carmo-Silva and Salvucci 2013). RCA functions in a hexameric ring structure (Portis 2003). Recent evidence indicates that the activity of RCA is largely modulated by its oligomeric state (Kuriata et al. 2014b; Henderson et al. 2013; Ning Zhang, Schürmann, and Portis 2001; Q. Wang et al. 2018). In the cell, RCA exists in three pools: inactive dimers, active hexamers, and inactive, thermostable higher-order aggregates. To function, RCA must form hexamers, catalyze hydrolysis of ATP (which can be coupled to rubisco activation), then dissociate into dimers, exchange ADP for ATP, and reform hexamers. Hexamer formation alone is not sufficient for activity; rather, the dynamic cycling of subunits between hexamer and dimer pools is required for activity. RCA transitions between hexamer and dimer pools involve the formation and breaking of inter-subunit contacts, the affinity and stability of which are determined by subunit concentration and modulated by the concentration of ATP, ADP and Mg^2+^.

ADP bound to individual subunits increases the rate of hexamer dissociation and results in a larger pool of inactive dimers, which may then form larger aggregates depending on magnesium concentration (Q. Wang et al. 2018; Kuriata et al. 2014a). Elevated ADP has been shown to improve the thermostability of RCA, likely by promoting thermoprotective aggregation (Henderson et al. 2013). RCA activity and aggregation is also sensitive to the concentration of Mg^2+^ in the stroma (Hazra et al. 2015; Barta et al. 2010), in part through its role in coordinating ATP binding, but also through a second, poorly characterized binding site (Kuriata et al. 2014a). Mg^2+^ promotes disassembly of higher-order oligomers and formation of hexamers, making subunits available to participate in the hexamer-dimer cycling that produces activity. Additionally, α and β RCA isoforms can form hetero-oligomeric complexes, and the thermostability and activity of the entire functional complex is thereby influenced by the conditions affecting both isoforms (Keown and Pearce 2014).

Efforts to find or create thermostable RCA isoforms have been advocated widely and have already shown moderate success (Davidi et al. 2020; Kurek et al. 2007; Scafaro, Bautsoens, et al. 2019; Qu, Mueller-Cajar, and Yamori 2023). To date, these efforts have not explicitly included possible regulatory effects of stromal conditions or post-translational modifications on either native or novel enzymes, making it likely that potential pitfalls and promising avenues for improvement have been missed. This is mostly due to an incomplete understanding of RCA regulation itself and in response to changes in its native environment: the dynamically changing chloroplast stroma. Toward this end, we examined whether CO_2_ assimilation is truly limited by RCA in the C_4_ food crops *Zea mays* (maize), *Sorghum bicolor* (sorghum) and in the common C_4_ model plant *Setaria viridis* (setaria). These species each possess α and β isoforms produced by dedicated genes. Intraspecific variation was assessed among several cultivars of maize, and interspecific variation was examined by comparison with sorghum and setaria. RCA isoform expression was determined in each species, and we examined how stromal conditions that are known to regulate RCA function may alter its activity during heat treatment. Our results expose potential avenues for improvement of RCA with the long-term goal to bioengineer crop species that maintain photosynthetic performance despite rising global temperature.

## 7 Materials and Methods

### 7.1 Plant Material

*S. viridis* A10 was a gift from the lab of Dr. Ivan Baxter at the Donald Danforth Plant Science Center in St. Louis. Seeds were sown 1cm deep in 18-well trays and thinned to one plant per cell. Plants were grown in the Washington University in St Louis greenhouse with 28°C/22°C temperatures and a 16-hour photoperiod. *Z. mays* B73 was obtained from the greenhouse at Washington University in St Louis. All other maize accessions were obtained from the USDA GRIN catalog. *S. bicolor* BTx623 was a gift from the lab of Dr. Malia Gehan at the Donald Danforth Plant Science Center. Sorghum and maize were planted 2cm deep in a 4-inch pot in a 50% mixture of Berger Bark Mix and Turface and grown in the Washington University in St Louis greenhouse with 28°C/22°C temperatures and a 16-hour photoperiod. Plants were grown to the two- or three-leaf stage, then transplanted into 6-inch pots in the same soil mix.

### 7.2 Heat Treatment

Greenhouse-grown plants were moved into a growth chamber of the appropriate temperature (42°C for most measurements.) Chambers were kept at 50% humidity and plants were maintained with the bottom of the pot in standing water to avoid drought stress. For 1-hour measurements, the leaf was acclimated in the gas exchange chamber of the LI-6400XT instrument with the instrument controlling the leaf temperature. For longer measurements, the gas exchange chamber was attached to the leaf twenty minutes prior to the measurement to allow stomata to equilibrate. For 48-hour measurements, plants were maintained at the same photoperiod as the greenhouse with 42°C day and 38°C night temperatures. All heat treatments were initiated and all gas exchange measurements were taken at least two hours from either end of the photoperiod.

### 7.3 Gas exchange measurements

Gas exchange measurements were performed on the most recently fully expanded leaf. The leaf measured was from the main stem of each plant, and all measurements were taken prior to panicle emergence. CO_2_ concentration within the gas exchange chamber was controlled at 400ppm. Gas exchange measurements were performed at saturating illumination of 1400 μmol m^-2^ s^-1^ (*S. viridis*) or 2000 μmol m^-2^s^-1^ (*Z. mays, S. bicolor*). Humidity was controlled to maintain stomatal conductance above 0.3 for all measurements. The same plants were measured at the 1, 6 and 48 hour time points whenever feasible.

A/Ci curve measurements were taken according to the standard method (Long and Bernacchi 2003) and were fitted using the Excel tool from (Zhou, Akçay, and Helliker 2018), which implements the model of (X. Yin et al. 2011) with the following modifications: changes to the constants found in cells P22, Q22, D126 and D127 (calculations of variables x_1_ and x_2_ for the electron transport limited state) were corrected to match the equations from (X. Yin et al. 2011), and α (the fraction of O_2_ evolution occurring in the bundle sheath) was assigned the value of 0 as PSII activity is negligible in bundle sheath cells in NADP-ME type monocot C_4_ photosynthesis (Von Caemmerer and Furbank 1999). For all measurements, the first measurement at 400μM CO_2_ was checked for stomatal limitation according to the equation in (Long and Bernacchi 2003) and any dataset in which stomatal conductance limited photosynthesis by more than 30% was discarded. This initial measurement was also compared to the A/Ci curve fit to determine the limitation to photosynthesis. Representative A/Ci curves and fits can be found in Figure S2, and all A/Ci curve fit variables may be seen in Table S1.

### 7.4 Tissue harvest and protein extraction

Leaf tissue from plants subjected to heat in growth chambers was harvested after 0, 1, 6 or 48 hours of heat exposure during the middle of the photoperiod, flash-frozen in liquid nitrogen, ground with mortar and pestle and pooled prior to storage at -80°C. All tissue samples contained leaves from at least 3 plants. Care was taken to exclude the ligule, the midrib, and leaves that were not yet fully expanded or which were senescent. RCA was extracted by ammonium sulfate precipitation according to the protocol of (Barta, Carmo-Silva, and Salvucci 2011a) with the modification that the 4.1M ammonium sulfate was adjusted to pH 4.6 instead of 7.0. Purified protein was resuspended in Leaf Resuspension Buffer (50 mM HEPES-KOH, pH 7.0, 2 mM MgCl_2_, 100 mM KCl, 5 mM dithiothreitol, 1 mM ATP, 1 mM PMSF, and 10 mM leupeptin). Aliquots were stored at -80°C until use. When compared to samples purified using the complete protocol of (Barta, Carmo-Silva, and Salvucci 2011a), samples using only the modified ammonium sulfate precipitation method were considerably cleaner and gave similar thermal curve results (Figure S5). Tissue was harvested from sorghum at 7 hours of heat treatment rather than 6 due to a gun-related lockdown at the ideal time of harvest.

### 7.5 Western blotting

Crude protein extracts from the first step of purification were resolved by SDS-PAGE on a 12.5% gel and transferred onto PVDF membrane using an Invitrogen Power Blotter semidry transfer apparatus according to the manufacturer’s instructions. 20μg of crude extract was used for maize, setaria and sorghum, and 5ug was used for reference species tobacco, Chlamydomonas, Arabidopsis and spinach. The membrane was blocked with 5% w/v nonfat dry milk in PBS supplemented with 0.1% v/v Tween-20 (PBST) with shaking at 4°C. Primary antibody (anti-RCA, rabbit, 1:10000 Agrisera) was added and incubated for one hour. The membrane was thoroughly washed with PBST and bound with secondary goat anti-rabbit-HRP (MilliPore) at 1:50,000 for 1 hour. The membrane was thoroughly washed and visualized with the FemtoGlow Western Plus chemiluminescent diagnostic kit (Michigan Diagnostics) according to the manufacturer’s instructions.

### 7.6 Recombinant protein production

The coding sequence of Mesembryanthemum crystallinum thioredoxin F was codon-optimized for expression in *Escherichia coli*, synthesized and cloned into pET28a with an N-terminal 6xHis tag. Transformed BL21-DE3 was grown to OD_600_ 0.6-0.8, induced with 1mM IPTG and incubated at 12°C overnight. TrxF was purified by Ni-NTA affinity chromatography in buffer containing 50mM HEPES pH 8.0, 0.5M NaCl, 5% glycerol, 10mM β-mercaptoethanol and 5-300mM imidazole, then dialyzed into the same buffer without imidazole and stored in aliquots at -80°C. TrxF activity was verified using the insulin reduction assay (Holmgren 1979).

### 7.7 Biochemical assays

RCA thermostability was measured with an NADH-linked ATPase enzymatic assay in microtiter plates based on the protocol of (Barta, Carmo-Silva, and Salvucci 2011b). Briefly, 20μL of purified RCA was aliquoted into 0.2mL tubes and incubated in a gradient thermocycler for 1 hour at temperatures ranging from 30-66°C and returned to ice until measurement, while a control aliquot was maintained on ice. RCA sample (5μL) was added to 120μL of reaction mixture in microtiter plates and the change in A_340_ was monitored for 10 minutes using a Tecan plate reader. ATPase activity was calculated as the initial slope of the line (change in absorbance per minute). T_50_ was calculated by fitting a 4-parameter logistic equation to the dataset using the calculator at (AAT BioQuest Inc, 2022), see example in Figure S4. Temperature response data that showed two distinct regions of response (sorghum and tobacco) were divided into two separate curves and each curve was fitted separately to produce two T_50_ values. The two plateaus are unlikely to represent a true response of RCA to heat. Previous work in spinach has shown that mixing of isoforms of different thermal stabilities confers an overall RCA response intermediate between the individual isoforms (Keown and Pearce 2014). Rather, the lower of the two T_50_ values is likely another ATPase present which was not fully removed by our purification technique. The portion of total ATPase activity due to the low-thermal-stability ATPase was calculated and subtracted from total ATPase activity results.

Response of ATPase activity to thioredoxin F, Mg^2+^ and ADP:ATP ratio was measured according to the method of (Chifflet et al. 1988; N Zhang and Portis 1999) because the presence of these small molecules interferes with the NADH-linked assay used to measure T_50_. Briefly, 40ug of purified enzyme was incubated in 100mM Tris-HCl pH 8.0 with 5mM DL Dithiothreitol with or without 2.6ug of TrxF as indicated for at least 20 minutes at room temperature to allow reduction of RCA. ATPase activity was then assayed in a total volume of 50uL of reaction buffer (50mM tricine pH 8, indicated concentration of MgCl_2_, 20mM KCl, 4mM total of ATP and ADP according to ratio) for 1 hour. Sodium dodecyl sulfate and ammonium molybdate reagent were added and allowed to react for 5 minutes. Color development was terminated by addition of sodium meta-arsenite reagent and A_850nm_ was read. Because the amount of ATP and the presence or absence of TrxF contributed different amounts to the background, no-RCA controls were performed for all treatments and the background was subtracted from all samples prior to statistical analysis. Measurements were normalized to ATP:ADP=1:0 without TrxF for all time points.

### 7.8 Statistical Analysis

Outliers were removed by Grubbs’ test, p<0.05, and significant differences between treatments were determined by T-test with Bonferroni correction. A two-way ANOVA was used to examine the interaction between ATP:ADP ratio and redox regulation.

## 8 Results

### 8.1 Response of CO2 Assimilation to heat

Sudden exposure to elevated temperatures results in a loss of CO_2_ assimilation capacity. We therefore first sought to obtain CO_2_ assimilation profiles from maize, sorghum and setaria exposed for 1h to a range of temperatures between 25 and 45°C (Figure S1 A-C). Peak assimilation occurred between 32-35°C for all species. When temperatures exceeded 40°C all plants displayed reduced capacity to assimilate CO_2_, with sorghum retaining a greater proportion of its peak activity at 42°C than maize B73 or setaria. At 45°C, assimilation had declined to below 20% of the maximum CO_2_ assimilation rate in all plants (Figure S1 A-C). In line with other studies, we therefore chose 42°C to study the impact of elevated temperatures on CO_2_ assimilation.

We next analyzed the capacity to acclimate the CO_2_ assimilatory capacity to elevated temperature in the different maize accessions using a time course, starting at 25°C (0h, prior to treatment), and sampling at 1 and 48 h after transition to 42°C. Maize B73 showed a significant reduction in CO_2_ assimilation capacity after 1h but managed to recover to levels similar to 25°C within 48h after transition to 42°C. MR15, MR25, MR26 and M162W were more resistant to the increase in temperature than B73, we found that 3 other cultivars shared the B73 response to sudden heat change, Tzi8, KI3 and Tx303. All four cultivars showed the same response, reduction after 1h, followed by a full recovery by the end of the study (Figure 1A). MR19 retained high CO_2_ assimilation rate after 1 hour but displayed a significantly reduced assimilation after 48 hours of heat exposure, indicating a failure to acclimate to elevated temperatures. This cultivar was chosen for further study because of its contrasting response to heat compared to B73. Chlorophyll fluorescence data for select maize cultivars is found in Supplemental Figure 3.

**Figure 1:**
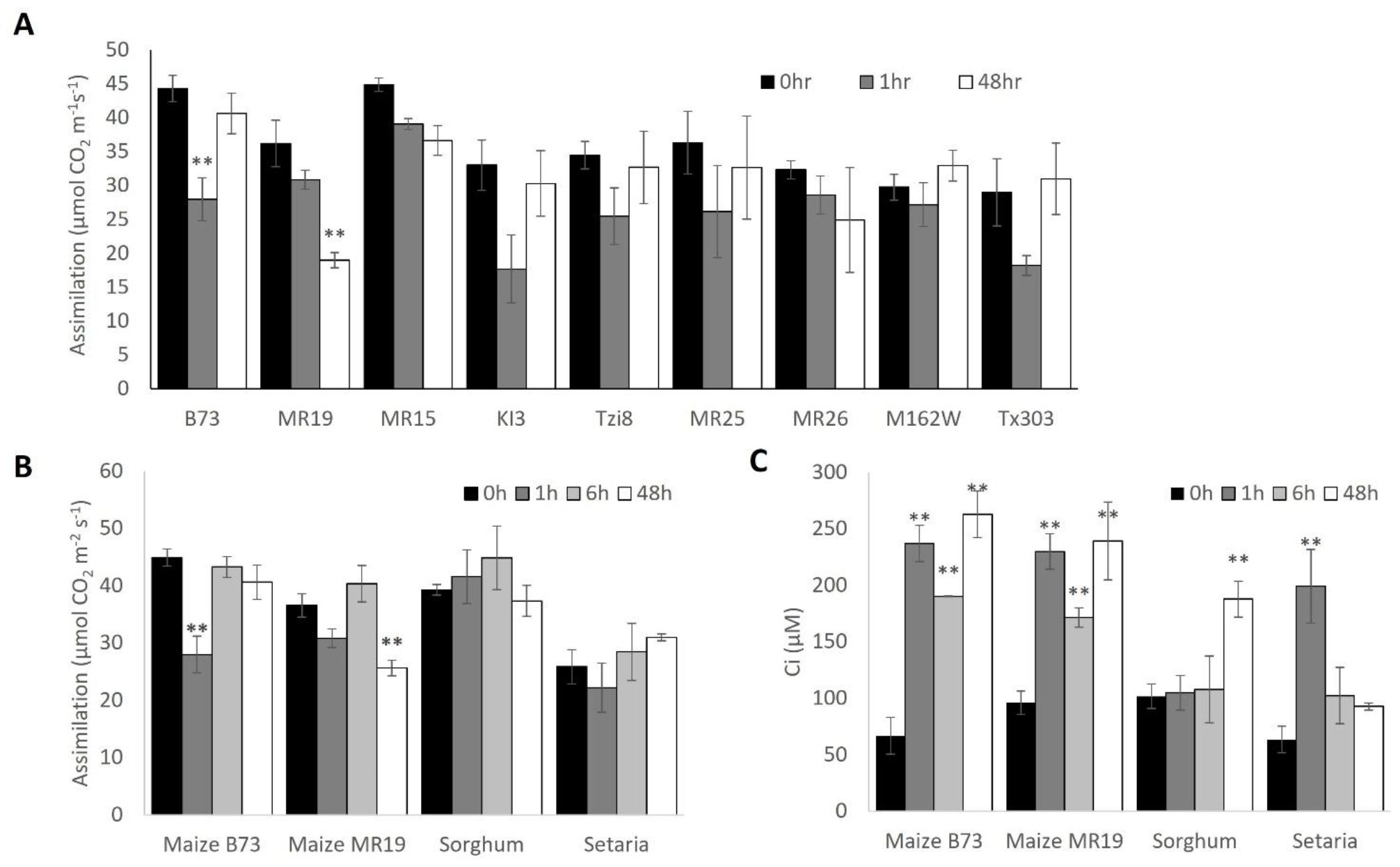
CO_2_ assimilation rates, intracellular CO2 levels (C_i_) and V_cmax_ during heat acclimation. **A**. Assimilation of various cultivars of maize. **B**. Assimilation of maize, sorghum and setaria. **C**. Ci of maize, sorghum and setaria. D. Vcmax of maize, sorghum and setaria. * p<0.05, ** p<0.01 vs. 0-hour control. Error bars are standard error of at least 3 biological replicates.

We next performed a more in-depth characterization of the heat response of CO_2_ assimilation for maize B73, maize MR19, *setaria viridis* A10 and *sorghum bicolor* BTx623. We measured both assimilation and response to CO_2_ (A/Ci) at 25°C (0h, prior to treatment), 1, 6 and 48 hours after transition to 42°C. The variance between individual replicates was lowest at 0 and 48 h, indicating that the plants successfully acclimated to the elevated temperatures by the end of the observation period and had reached a stable state. Variance between individual plants was higher during acclimation, and most prominently after 1h at 42°C, indicating some differences in timing and responsiveness between individual plants, despite all being exposed to the temperature conditions at the same time. Sorghum and setaria were quite resilient to sublethal heat, showing consistent CO_2_ assimilation throughout the course of the acclimation experiment.

Intercellular CO_2_ concentration (C_i_) was measured for all species at normal and elevated temperature at 400ppm CO_2_ (Figure 1C). All species exhibited elevated C_i_ at high temperatures. In both maize cultivars, the Ci more than doubled within 1 hour and remained elevated. In sorghum, C_i_ remained low until the 48-hour time point. In setaria, the increase in C_i_ was transient, and C_i_ returned to pre-treatment values within 6 hours. Stomatal conductance in all plants increased during acclimation and paralleled the trends seen in C_i_ (Supplemental Figure 1D).

### 8.2 Limitations to Photosynthesis

The response of assimilation to changes in intercellular CO_2_ concentration (C_i_) can be measured and mathematically fitted to find values of photosynthetic parameters and to determine which factor is limiting photosynthesis. At low C_i_, photosynthesis is limited primarily by the speed of the carboxylation reaction carried out by rubisco. At higher C_i_, photosynthesis is primarily limited by the ability of the CBB cycle to regenerate the RuBP substrate of rubisco, which in turn is limited by the rate of electron transport to regenerate reducing equivalents for this process (Long and Bernacchi 2003). Additionally, in C_4_ plants which use phosphoenolpyruvate (PEP) to transport CO_2_ from the mesophyll cells to the bundle sheath cells, the speed of PEP carboxylation or regeneration also affect CO_2_ assimilation.

To determine the limitation to CO2 assimilation at atmospheric CO2 levels (400ppm), we measured CO_2_ assimilation dependent on C_i_ (A/Ci curves) in triplicate for each of the four plants after 0, 1, 6 and 48 hours of heat exposure and fit the measurements for each plant to a model of C_4_ photosynthesis, as described in detail in (Zhou, Akçay, and Helliker 2018) (Figure S2 and Table S1). At ambient temperatures both maize cultivars were strictly limited by the carboxylation reactions, while one sorghum plant was limited by RuBP regeneration rather than RuBP carboxylation. All setaria cultivars were limited by RuBP carboxylation, but one setaria plant was concurrently limited by PEP regeneration. After transition to elevated temperatures (1 and 6h) all plants were in strictly carboxylation limited both for RuBP and PEP. Interestingly, all sorghum plants at 48h were limited by RuBP regeneration, while both setaria and the two maize cultivars were still limited by carboxylation. The A/Ci curve fits additionally provided five photosynthetic parameters, including V_cmax_, the maximum carboxylation rate of rubisco (Figure 1D). The overall pattern looks largely similar to CO_2_ assimilation rates. V_cmax_ was significantly decreased in maize cultivar MR19 at the 48h time-point, when it also exhibited compromised CO_2_ assimilation.

Taken together, RuBP carboxylation was the major limiting factor to C_4_ photosynthesis at high temperature. This is consistent with RCA failing to maintain rubisco in a catalytically competent state. Only in sorghum we found a shift away from the carboxylation reaction towards the regeneration of the RuBP to contribute to photosynthetic limitation, but only after extended periods of exposure.

### 8.3 RCA expression and proteoform composition in response to heat

Heat inactivation of RCA appears to play a major role in maintaining photosynthetic performance during heat acclimation in all plants investigated in this study. Next, we therefore shifted our attention to RCA directly and determined the abundance and proteoform composition in the two maize cultivars, sorghum and setaria. Immunodetections were performed on crude leaf tissue extracts before and after heat treatment, again time-dependently, with RCA-specific antiserum (Figure 2). Maize cultivar B73 was the only plant in this study that showed little expression of RCA at ambient temperatures, which may account for the large reduction to CO_2_ assimilation early after transfer to elevated temperatures (Figure 2, Figure 1B). RCA abundance increased after transfer to higher temperatures in maize and sorghum, evident already after 1h of heat treatment and peaking around the 6h time point. Setaria did not exhibit increased RCA abundance. For all plants we found the expression of at least 3 RCA proteoforms. In order to determine the identity of the different proteoforms (α, β, etc.) we compared our results with the RCA composition in model organisms that had been described previously in the literature (Figure 2). Arabidopsis and spinach samples contain an α isoform at 45kDa, β at 43kDa and a proteolytic product of β at 41kDa, with Arabidopsis proteins running slightly higher than their spinach equivalent. Tobacco and Chlamydomonas only express the β isoform, and in our detections both species lack the higher molecular weight bands corresponding to the α isoform. The lower band intensity for the alga *Chlamydomonas reinhardtii* could indicate a generally lower RCA abundance, an adaptation to the presence of a carbon concentrating mechanism (CCM), but more likely may indicate lower antibody affinity for this more evolutionary distant homolog. While Chlamydomonas shows only a single proteoform, tobacco displays several proteoforms of β, including a band at a slightly lower molecular weight of about 39kDa (Figure 2, see also complete blots in Figure S3). In both maize cultivars and in sorghum we found an additional higher MW band that appeared only after heat treatment, most prominently around the 6h mark. Although the maize α isoform is known to be phosphorylated at threonine 78, we hesitate to positively identify this band because its size shift seems larger than could be explained by a single phosphate. Setaria does not contain the threonine at this position, and we did not observe a higher molecular weight form of α in our immunodetections. Other changes in proteoform abundance were specific to the species or cultivar. In setaria the α isoform is depleted within 1h of heat treatment and does not recover during the observation period. The 41kDa β proteoform is transiently enriched in response to heat. In maize MR19 there is a transient decrease in the unprocessed α isoform, after 1h of heat exposure. Sorghum shows the least amount of change in protein composition overall outside of the appearance of the phosphorylated product.

**Figure 2:**
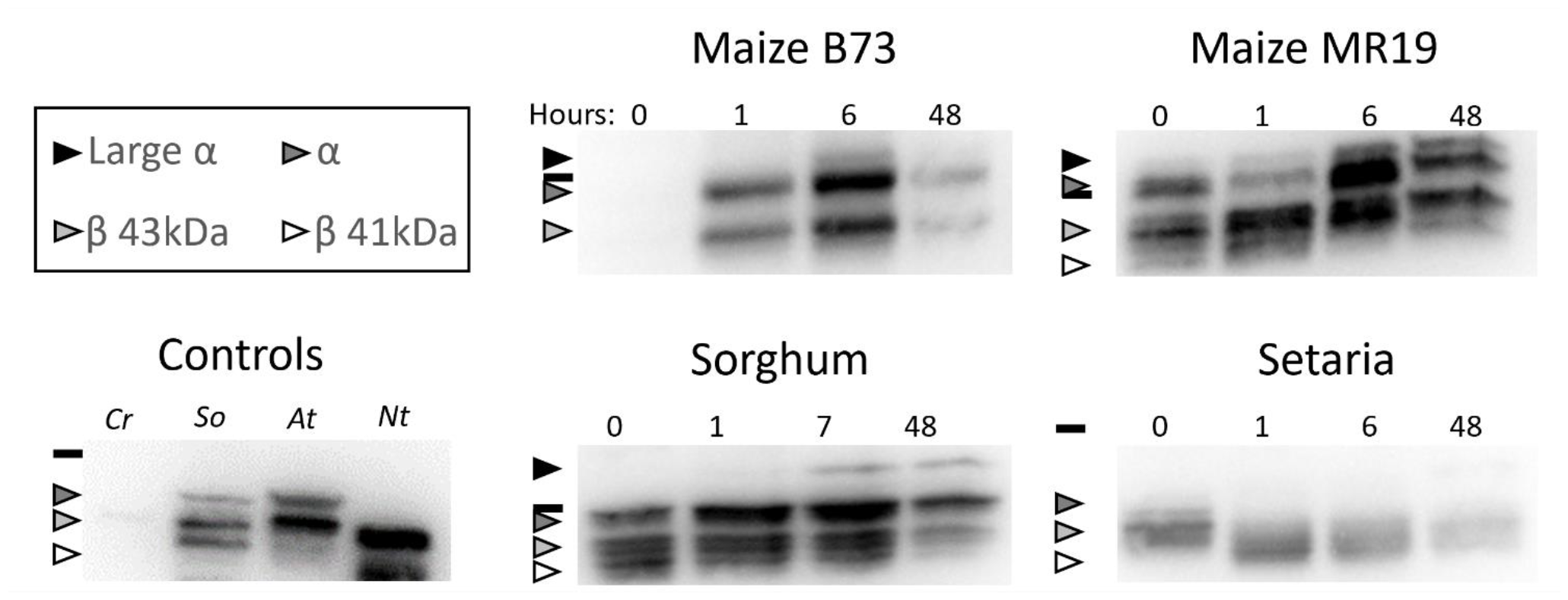
RCA abundance and proteoform distribution during heat acclimation. Immunodetections were done on 20μg of crude protein extract for C_4_ species and 5μg of crude protein extract for controls. Cr *C. reinhardtii*, So *S. oleracea*, At *A. thaliana*, Nt *N. tabacum*. Times are indicated for each sample in hours. Complete blots may be viewed in the supplement. Black line indicates 37kDa marker; proteoforms are indicated by shaded arrows according to the legend

Taken together, induced RCA expression and presence of the higher molecular weight α proteoform appear to be critical components in maize and sorghum, but are not involved in heat acclimation in setaria. Changes in the proteolytic processing of the β isoform was only observed in setaria in response to elevated temperatures.

### 8.4 RCA ATPase activity in C_4_ plants at elevated temperatures

We next characterized RCA’s enzymatic ATPase activity in different settings relevant for heat acclimation. Recombinantly produced proteins are commonly used for biochemical assays, but due to the complexity of RCA proteoforms and importance of post-translational modifications, we decided to purify RCA directly from plant tissue.

First, we examined how RCA’s ATPase activity is affected by exposure to elevated temperatures. We normalized the ATPase activity either per ug isolated protein (black bars), to capture the intrinsic activity of RCA itself, which can change due to modifications and proteoform composition changing upon heat exposure, or to total leaf content (white bars), allowing to assess the total capacity of RCA for rubisco reactivation, including adjustments made to total RCA protein content. In maize cultivar MR19, which fails to acclimate to heat (Figure 1), total RCA activity and intrinsic activity are closely linked and were reduced dramatically directly after exposure to elevated temperatures (1h time point, Figure 3B). RCA activity subsequently only recovered slightly, resulting in only ∼70% of ambient total RCA activity retained at 42°C. In the other maize cultivar, B73, total RCA activity increased substantially, by ∼ 7-fold, at 42°C, peaking at 6h after heat exposure (Figure 3A). This increase was solely driven by the increase in RCA protein abundance (Figure 2), since intrinsic RCA activity (per μg protein) was reduced immediately after heat exposure and only recovered slightly by 48h, in a similar profile than maize MR19 (Figure 3A,B). Both sorghum and setaria showed a close connection of intrinsic and total RCA ATPase activity, but, while sorghum peaked at the 1h mark, it took setaria much longer to increase its RCA activity. By the end of the experiment, intrinsic RCA activity was dramatically increased in setaria (∼5 fold), while ATPase activity in sorghum was fully attenuated back to initial levels (Figure 3 C,D).

**Figure 3.**
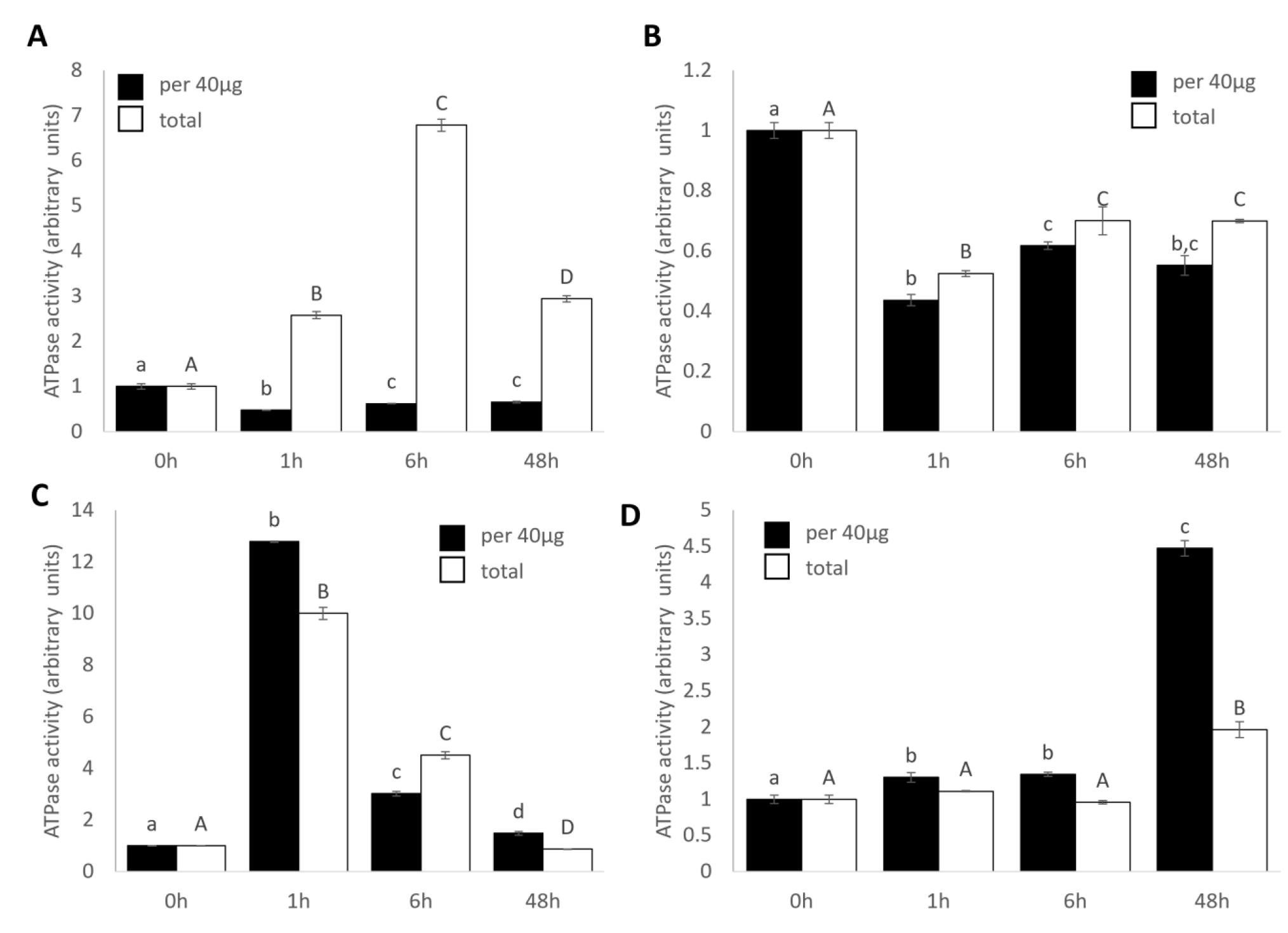
ATPase activity of RCA. **A**. maize B73 **B**. maize MR19 **C**. sorghum **D**. setaria. Letters denote significance within data series of p<0.05. Error bars are standard deviation of n=5

**Figure 4.**
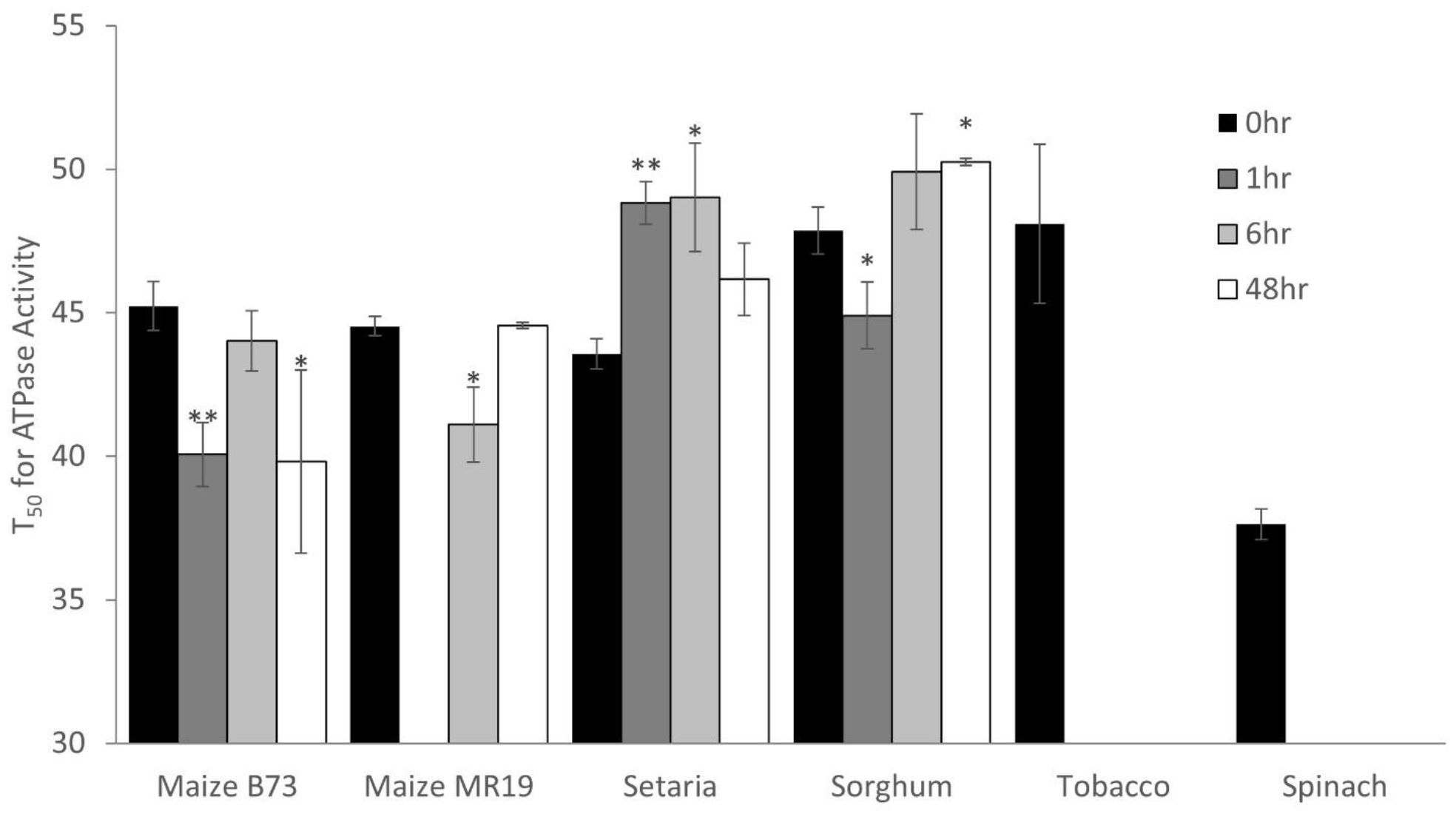
temperature sensitivity of ATPase activity. * p<0.05, ** p<0.01 relative to 0 hours, error bars are standard deviation of n=3 technical replicates

Taken together, RCA ATPase activity in maize was reduced early after exposure to elevated temperatures, which appears to be compensated by increasing overall RCA abundance in thermotolerant cultivars. Sorghum and setaria both manage to increase intrinsic ATPase activity in response to heat to maintain their CO_2_ fixation capacity, but are on very different timelines.

### 8.5 RCA thermostability

Our data indicate that different plant species (and even different cultivars of the same species) show flexibility in how RCA abundance and activity change during heat acclimation. We next sought to determine how these changes affect RCA thermostability. We measured RCA thermostability by pre-incubating purified enzyme for 1 hour at temperatures ranging from 30°C to 66°C, then measuring activity using a spectroscopic ATPase assay based on NADH consumption. The resulting heat response curves were fitted to determine the temperature at which half of the activity was lost (T_50_). As expected, spinach RCA had a lower T_50_ than tobacco RCA (37.6°C vs. 48.0°C). The thermostability of RCA extracted from both maize cultivars varied with time but never increased beyond the 0-hour control. In contrast, both Setaria and sorghum RCA displayed increased T_50_ at various times, with Setaria increasing from 43.6°C (0-hour) to 49.0°C (6 hours) and sorghum increasing from 47.9°C (0 hour) to 50.3°C (48 hours).

### 8.6 Effect of Mg^2+^ on RCA ATPase activity

The availability of magnesium can affect the activity of many enzymes, especially ATPases. We therefore studied the responsiveness of RCA’s ATPase activity to changes in Mg^2+^ concentration. At ambient temperatures, all other plants show some responsiveness to changes in Mg^2+^ with the exception of maize MR19 which didn’t show any difference and fails to acclimate to elevated temperatures, (Figure 5). Most proteins were significantly more active with more Mg^2+^ in the buffer (Figure 5). These increases are consistent with what had been previously reported for tobacco (Hazra et al. 2015). From the C_4_ plants analyzed here only sorghum was stimulated by Mg^2+^; supplementing 10 mM Mg^2+^ increased RCA activity by 43% (Figure 5D). To our surprise, both maize B73 and setaria showed inhibition by increased Mg^2+^ concentrations (Figure 5B,E). For maize, the inhibition was dramatic (34% activity inhibited by 10mM Mg^2+^, p<0.01), setaria showed less of an effect. We also analyzed Mg^2+^ responsiveness following heat treatment, to see if any of the modifications to the proteoform composition or post-translational modifications affected the regulation by Mg^2+^. In sorghum, responsiveness to Mg^2+^ was transiently dampened, to below < 10% increase at 1 and 7 hours of heat treatment. B73 RCA, which was most strongly inhibited by Mg^2+^ at ambient temperatures, lost the Mg^2+^-dependent inhibition gradually after heat exposure. For setaria, the inhibition was less pronounced (9% at 10mM Mg^2+^, not significant) at ambient temperatures and only rose to the level of significance after 48 hours of heat treatment (16% inhibition at 10mM Mg^2+^, p<0.01).

**Figure 5.**
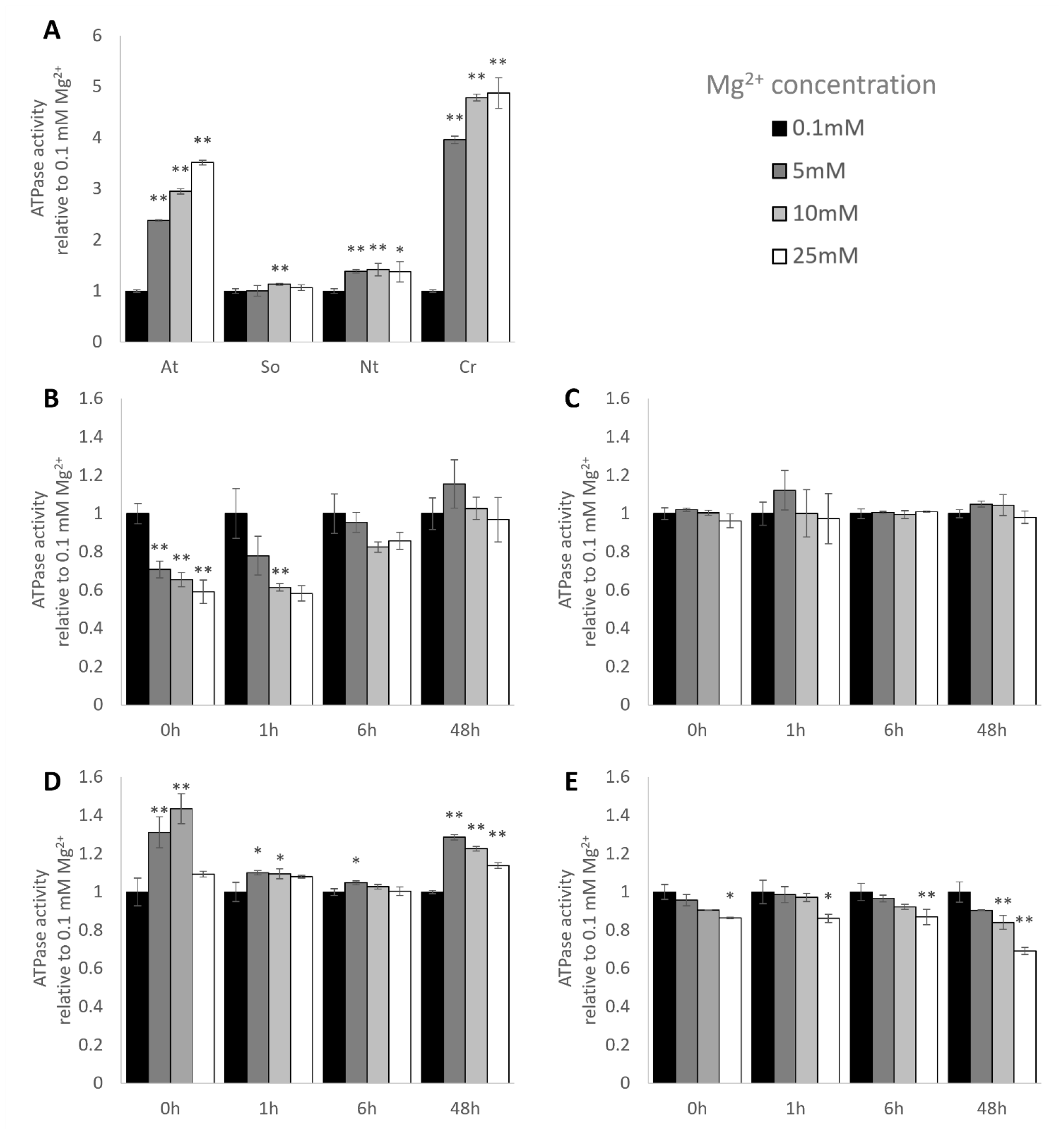
response of ATPase activity to changes in Mg^2+^ concentration. All measurements performed with ATP:ADP=4:0. **A**. Controls **B**. Maize B73 **C**. maize MR19 **D**. sorghum **E**. setaria. * p<0.05, ** p<0.01 relative to 0.1mM Mg^2+^, error bars are standard deviation of n=5.

### 8.7 Effect of ATP/ADP ratio on RCA ATPase activity

Previous work has established that the ATPase activity of RCA is inhibited by increasing concentrations of ADP (Ning Zhang, Schürmann, and Portis 2001). The ATP:ADP ratio is a good indicator of the energy status of the chloroplast: a high ATP:ADP ratio is usually seen in plants with a highly active photosynthetic electron transfer chain, while high amounts of ADP indicate an inadequate supply of ATP for chloroplast energy-dependent metabolic reactions, most prominently carbon fixation via rubisco. We were able to replicate the ADP-dependent inhibition of RCA’s ATPase activity with RCA purified from Arabidopsis, spinach, tobacco and Chlamydomonas, which were most affected by high amounts of ADP in that order (Figure 6A). At ambient temperatures, ATPase activity in RCA from setaria was similarly inhibited by ADP. While maize B73 was not affected by the ATP:ADP ratio at all, sorghum and maize MR19 showed increased ATPase activity with a higher ADP:ATP ratio (Figure 6). In response to elevated temperatures, maize B73 eventually showed slight stimulation by increased levels of ADP. Maize MR19 remained stimulated by ADP at all times (Figure 6). Sorghum RCA lost its sensitivity to ADP immediately upon heat exposure. Setaria also lost ADP sensitivity, but only after 48 hours of heat. Taken together, setaria plants were the only ones in our set of C_4_ plants that showed classic ADP-dependent inhibition of ATPase activity, all other species showed some amount of stimulation by increased ADP amounts.

**Figure 6.**
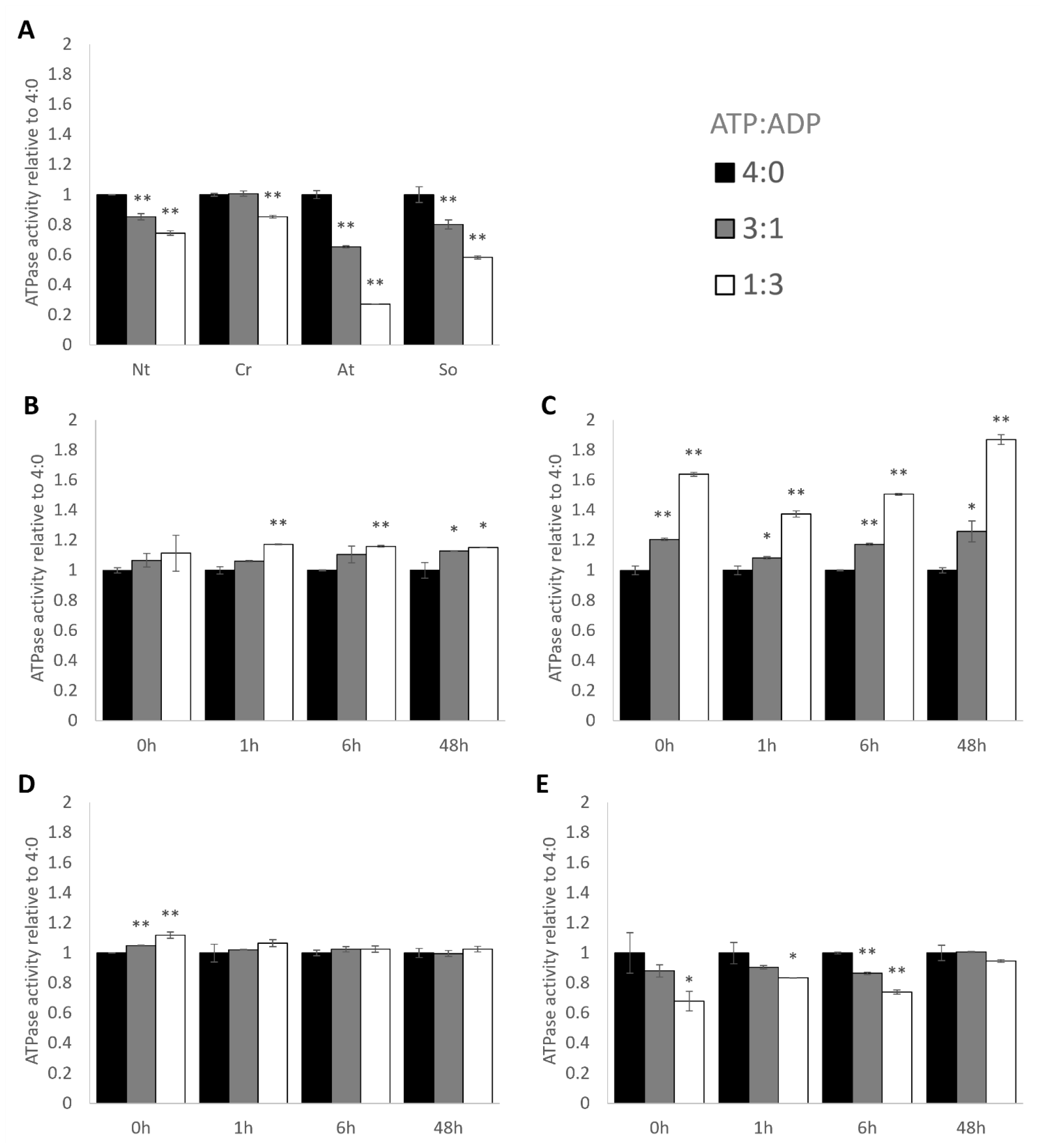
response of ATPase activity to changes in ATP:ADP ratio. All measurements performed with 10mM Mg^2+^. **A**. Controls **B**. maize B73 **C**. maize MR19 **D**. sorghum **E**. setaria * p<0.05, ** p<0.01 relative to 4:0 ratio, error bars are standard deviation of n=5

### 8.8 Redox sensitivity

TrxF-mediated redox regulation of the disulfide bond in the C-terminal domain of the α isoform of RCA is known to be important in C_3_ plants (Ning Zhang, Schürmann, and Portis 2001; Ning Zhang et al. 2002). However, in NADP-ME-type C_4_ plants such as maize, sorghum and setaria, NADPH reducing equivalents are imported into bundle sheath cells from mesophyll cells, and electron transport in BSCs is therefore specialized for ATP production via cyclic electron flow rather than for producing reducing equivalents. The redox state of BSCs has not been well characterized, and the extent of TrxF-mediated redox regulation is currently unknown. Nevertheless, we examined this response in our C_4_ plants. Maize RCA was slightly stimulated by TrxF-mediated reduction, and sorghum and setaria were insensitive (Supplemental Figure 4).

## 9 Discussion

Analyzing gas exchange measurements, enzymatic activities, protein abundance, modification and proteoform composition allowed us to compare the properties of different RCA proteins across C_4_ monocot plants which paves the way in our understanding of the acclimation strategies that these plants employ to maintain CO_2_ assimilation at elevated temperatures. We found that in all plants, similar to earlier reports, CO_2_ assimilation at high temperature was limited by RuBP carboxylation, and therefore likely by RCA activity. There were, however, substantial differences in the way these crops employed the various levels of RCA regulation to maintain photosynthetic carbon fixation.

### 9.1 Species-specific strategies for high temperature acclimation

Setaria, maize and sorghum assimilated CO_2_ efficiently over a range of temperatures up to 40 °C, where we first found CO_2_ assimilation rates starting to decline. At 42°C heat, all plants maintained their carbon fixation capacity after 48 hours of exposure, with compensating changes apparent both at the physiological level (e.g. stomatal opening) and molecular level (e.g. protein abundance and proteoform composition). Notably, maize cultivar MR19 failed to acclimate to high temperature despite physiological and molecular changes and did not maintain its initial CO_2_ assimilation rate after 48 hours. Altered proteoform composition translated to changes in RCA ATPase activity, especially with regards to response to local environmental stimuli (e.g. temperature, Mg^2+^ and ADP/ATP ratio). However, the timing, magnitude, and direction of all of these changes differed substantially between the different species.

#### 9.1.1 Maize

There was a wide range of CO_2_ assimilation responses observed at 42°C across multiple maize cultivars. While several maize cultivars showed no decline in CO_2_ assimilation, even early during heat acclimation, maize B73 transiently decreased CO_2_ assimilation and MR19 failed to acclimate after extended exposure to heat. In both cultivars, rapid physiological changes occurred within the first hour of heat exposure: increases in stomatal conductance and a corresponding increase in C_i_. Both cultivars remained limited by RuBP carboxylation, and there was no change to V_cmax_ throughout the duration of heat treatment. During acclimation, there were dynamic biochemical changes in B73: a rapid but transient increase in RCA abundance, appearance of a higher molecular weight proteoform of α, rapid gain of stimulation by ADP, gradual loss of inhibition by Mg^2+^, and a slow increase in q_N_ (Supplemental Figure 3). While both setaria and sorghum manage to increase their intrinsic RCA ATPase activity as well as the thermostability (T_50_) in response to heat, both maize cultivars fail to do either, and even showed reduced ATPase activity after heat exposure. B73 however compensates by dramatically increasing the abundance of RCA, managing an overall increase in RCA ATPase activity, which does not occur in MR19. In contrast, the biochemistry of MR19 was more similar to the B73 acclimation endpoint at all time-points: even at the 25°C 0-hour measurement, MR19 displayed high RCA abundance, ADP sensitivity, Mg^2+^ insensitivity and high q_N_. The appearance of the higher MW proteoform of α was a transient event in B73, while it can be observed in MR19 at all time points, even at ambient temperatures (time point 0h). It appears that even at the greenhouse condition (28°C), MR19 constitutively activates the biochemical program of RCA heat acclimation but cannot sufficiently elevate its RCA activity when further challenged at 42°C heat.

#### 9.1.2 Sorghum

In sorghum, stomatal conductance and C_i_ changed only after prolonged heat treatment. However, rapid biochemical changes occurred. An immediate, transient increase in RCA abundance was accompanied by a suppression of stimulation by both ADP and Mg^2+^. After 7 hours of heat treatment, the higher molecular weight α proteoform appeared and remained even when overall RCA abundance decreased at 48 hours. The occurrence of the higher larger RCA coincided with an increase in thermostability, which was not observed in maize. Mg^2+^ sensitivity, but not ADP sensitivity, returned to initial levels by 48 hours. Unique to sorghum out of all plants examined was that, by the end of acclimation, RuBP carboxylation was no longer limiting for CO_2_ assimilation. It is noteworthy that in both maize and sorghum, although the increased abundance of RCA was transient, the changes in biochemical properties were sustained. This suggests that these biochemical properties derive from the altered composition of RCA proteoforms, such as might be produced by post-translational modifications. Further study into the exact identities and properties of these proteoforms should be pursued.

#### 9.1.3 Setaria

While maize and sorghum both exhibit an immediate increase in RCA abundance and rapid changes to RCAs’ enzymatic properties, setaria RCA protein abundance never increased, and enzymatic changes took at least 6 hours to manifest. Instead, setaria appears to execute an immediate but transient C_i_ increase mediated by marginal changes in stomatal conductance. Although the abundance of RCA appears unchanged throughout the time course, the proteoform composition of the lower molecular weight bands is shifted early, coinciding with a small but a significant, temporary increase in RCA T_50_. At 48 hours, setaria displayed elevated per-molecule ATPase activity, loss of ADP inhibition and slight increase in Mg^2+^ sensitivity. Unique among the C_4_ species studied, setaria is ADP-inhibited, matching the profiles of C_3_ species. As with the other species, Vcmax remained unchanged throughout heating.

### 9.2 Small molecule regulation may control oligomeric state

The genetically programmed response to a rise in higher temperatures seems to impact C4 plants vastly different as compared to C_3_ plants or algae when it comes to changes in the biochemical inventory of the chloroplast stroma. For example, in C_3_ plants and algae, Mg^2+^ stimulated ATPase activity in a concentration-dependent manner, but inhibited activity in both maize and setaria. Similarly, high ADP:ATP environments were inhibitory to control plant RCAs but stimulated those from maize and sorghum. We hypothesize that these results point to differences in the oligomerization of C_4_ RCA as described in the following. Experimental evidence has resulted in a model of three oligomeric states which coexist within the chloroplast stroma. Inactive dimers assemble into active hexamers, which can then form higher-order inactive aggregates. There is an optimal equilibrium ratio between hexamer and dimer states which produces the highest activity, and deviations from that optimum reduce RCA activity. ADP promotes the formation of dimers and, at high RCA concentrations, aggregation. Mg^2+^ promotes disassembly of aggregates and formation of hexamers. For this reason, in our preparations of C_3_ plants, Mg^2+^ increases the ATPase activity of RCA by liberating subunits from aggregates and increasing the pool of actively cycling RCA, while ADP decreases activity by shifting the hexamer-dimer equilibrium toward a suboptimal, dimer-dominated state (Figure 7, top row). Kuriata et al 2014 demonstrated that the affinities for hexamer formation and for aggregation are at least partially independent from each other insofar as ATP, ADP and Mg^2+^ produced different effects on each. It has also been shown that in a mix of proteoforms, the properties of each proteoform contribute to determine the oligomerization state of the entire pool.

**Figure 7.**
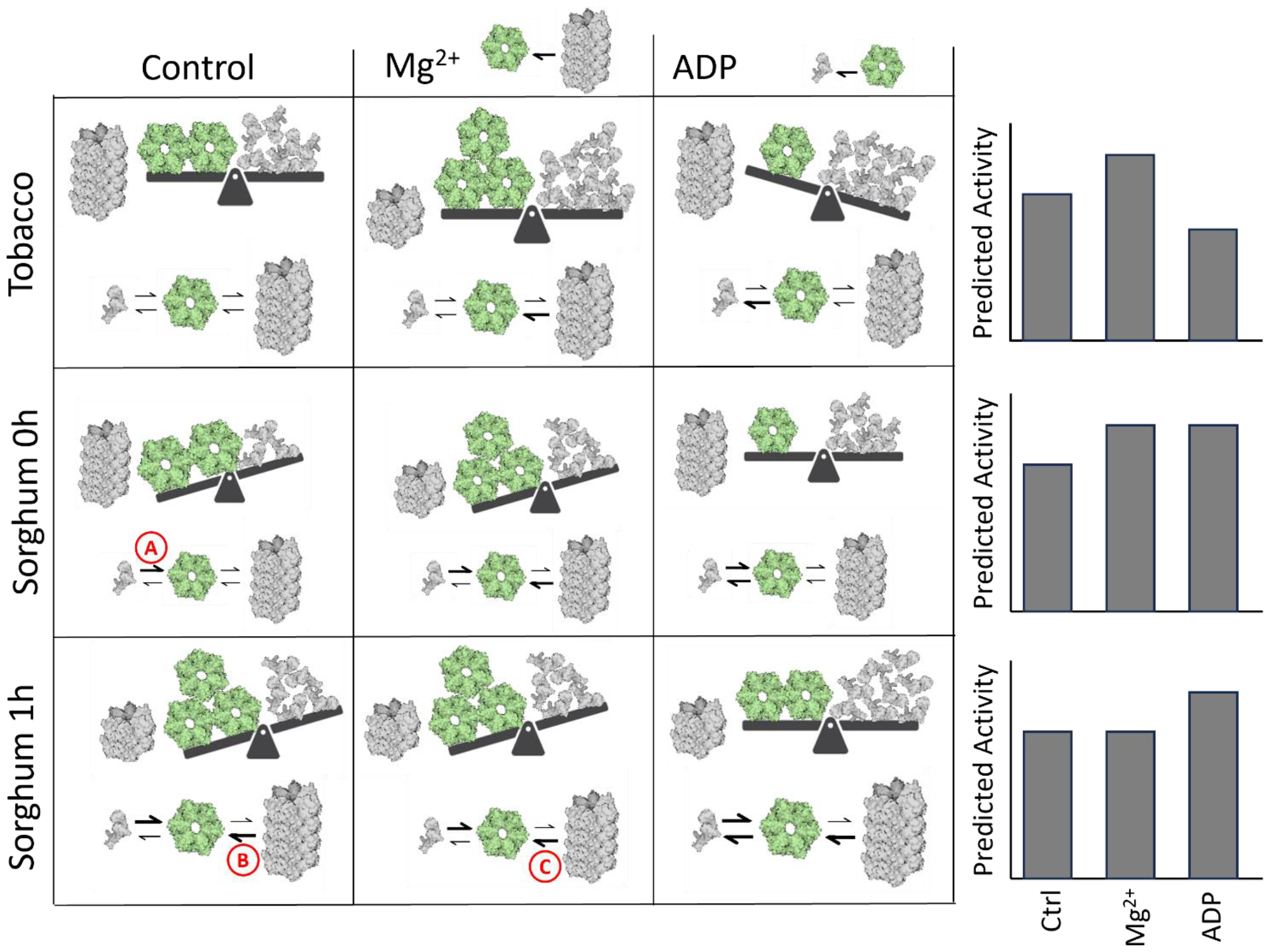
a proposed mechanism for heat-mediated changes in RCA ATPase activity and small molecule responses. Imbalance between dimers and hexamers always reduces activity. See further description in the text of the Discussion.

We hypothesize that the changes in proteoform composition seen after heat treatment shift the relative affinities for hexamer formation vs. aggregation. Imagine that the hexamer formation affinity in sorghum RCA is higher than that of tobacco (Figure 7, arrow labeled A). Thus, at the 0h timepoint, subunits were primarily in the aggregated state and in the hexamer state. Addition of Mg^2+^ would stimulate activity by releasing more subunits from aggregates to participate in the dimer-hexamer cycle, while addition of ADP would increase activity by rebalancing the cycle toward the underrepresented dimer state (Figure 7, second row). After 1 hour of heat, changes in proteoform composition could reduce the subunit aggregation affinities (Figure 7, arrow labeled B). Because proteoforms interact to determine the overall phenotype, changes to a small proportion of subunits could destabilize all aggregates, shifting a large proportion of subunits from aggregates into the actively cycling pool. In this way, for example, a loss of Mg^2+^ sensitivity by sorghum isoforms at 1 hour (Figure 5) could indicate that the proteoform affinity has changed such that higher-order aggregates have already been destabilized and more subunits cannot be liberated into the active pool by adding Mg^2+^ (Figure 7, arrow labeled C) When a larger proportion of subunits is present in the actively cycling pool than in the inactive aggregate, the activity per unit protein would increase, as observed in Figure 3. Similar explanations based on subunit affinities can be made for the activities of maize and setaria, and experiments like those performed by Kuriata et al 2014 should be used to quantitatively test this proposed mechanism.

## 10 Conclusions

Heat acclimation is a dynamic process in which plants modulate their stromal environment and RCA proteoform abundance, accompanied by physiological changes like stomatal conductance affecting intercellular CO_2_ concentration. All these parameters must be considered to obtain a holistic understanding of the mechanisms contributing to thermal limitation in each plant at each time point.

One example is the observation that increased RCA activity in setaria and sorghum is not fully explained by increased RCA abundance. This implies that while overexpressing RCA may improve heat tolerant carbon assimilation in some plant species (Qu et al. 2021), plants evolved many more mechanisms for promoting RCA activity without overexpression, some of which are showcased in this work. Furthermore, the thermostability of the enzyme itself is not predictive of its activity, so attempts to engineer a more thermostable protein may not improve activity *in planta*. Rather, we have shown that RCA activity is regulated in response to heat by a combination of post-translational modifications and by small molecules, which likely affect the oligomerization state of RCA. The magnitude and direction of these effects is species-specific and dependent on the duration of the heat treatment. Thus, we caution that observations regarding the heat acclimation strategy in the model species setaria cannot reliably be transferred and used to bioengineer other crop plants. Nor can the effects of the biochemical environment be safely extrapolated even among cultivars of the same species. Future work must therefore be specific to the species and cultivar.

We suggest that future studies should focus on determining the identities of heat tolerant RCA proteoforms that are biosynthesized during plant heat acclimation, and on potential changes in abundance of hexamers and aggregates during heat acclimation. It is clear that *in vitro* assays will not be able to fully capture the biochemical complexity of the chloroplast stroma as well as the proteoform diversity that we observed. A more mechanistic understanding of the molecular events during heat acclimation *in vivo* will likely generate actionable predictions that could improve assimilation in these plants during heat exposure.

## 3 Author contributions

Conceptualization: SCS. Methodology: SCS, LNA, AC, EJ, AS, LW, AA, JJ. Validation: SCS, LNA, AC, EJ, LW, AA. Investigation: SCS, LNA. Analysis: SCS, LNA, AC, EJ, AS, LW. Writing: SCS, LNA. Review and editing: SCS, LNA, AC, EJ, AS, LW, AA, DS, JJ. Funding: JJ, DS, SCS. Supervision: JJ, DS.

## 4 Data Availability

All data is contained within this paper and its supporting material, and raw data files are available upon request to the corresponding author at.

## 11 Acknowledgments

We thank Ru Zhang for the initial inspiration of this work, Tom Sharkey, Oliver Mueller-Cajar, Stefan Schmollinger and Kevin Wyss for feedback on the manuscript, Leonardo Chavez for assistance with the LI-6400XT machine and Mike Dyers for assistance with plant growth. This work was supported by the Amgen summer scholar program (LNA) and an NSF Postdoctoral Research Fellowship in Biology #1907288 (SCS).

