## Supplementary Material for "C_4_ Grasses Employ Distinct Strategies to Acclimate Rubisco Activase to Heat Stress"

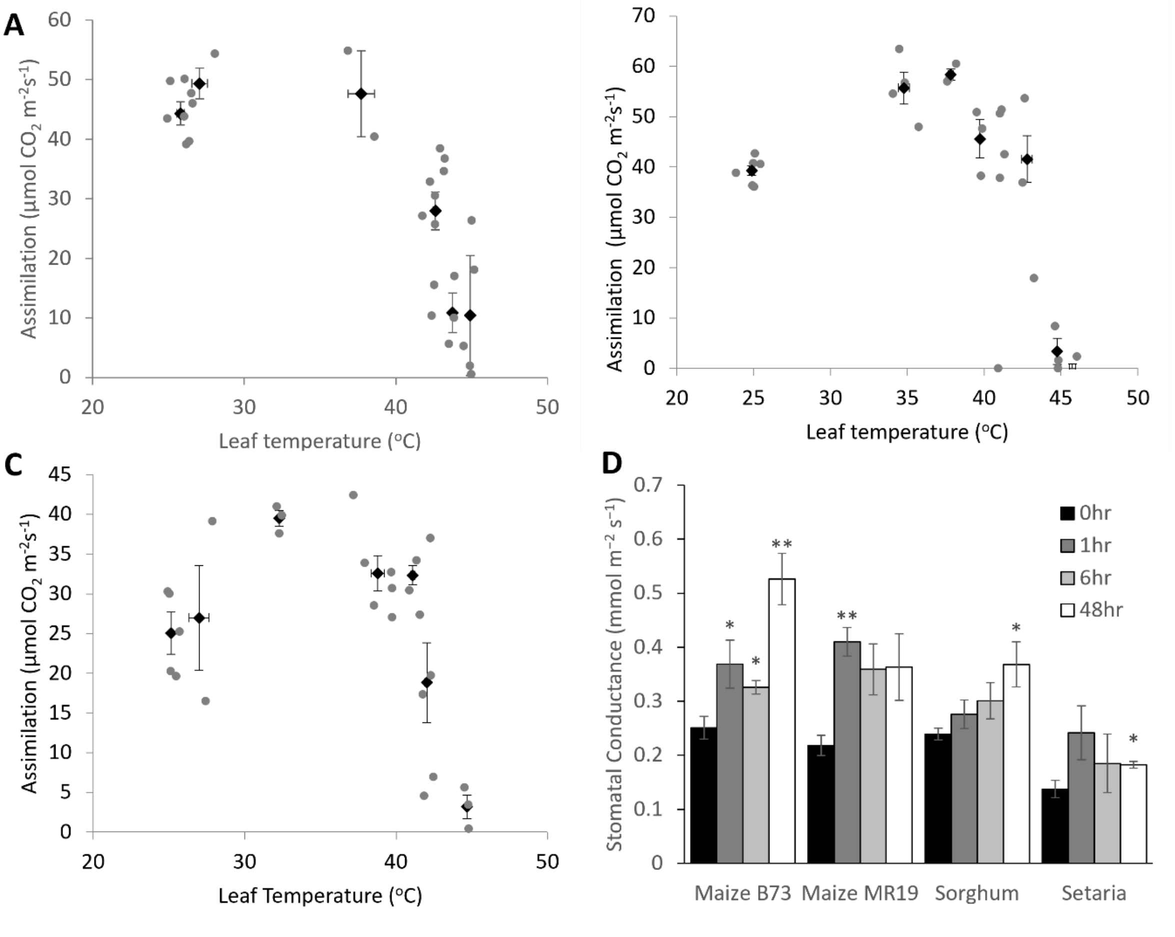


**Figure S1**: Additional gas exchange data. **A-C.** assimilation after 1 hour treatment at various temperatures. **A.** Maize B73. **B.** Sorghum. **C.** Setaria. Grey points are individual plants, dark points and error bars are mean and standard error of at least 3 measurements. **D.** Stomatal conductance of maize, sorghum and setaria. * $p<0.05$, ** $p<0.01$ vs. 0-hour control. Error bars are standard error of at least 3 biological replicates.

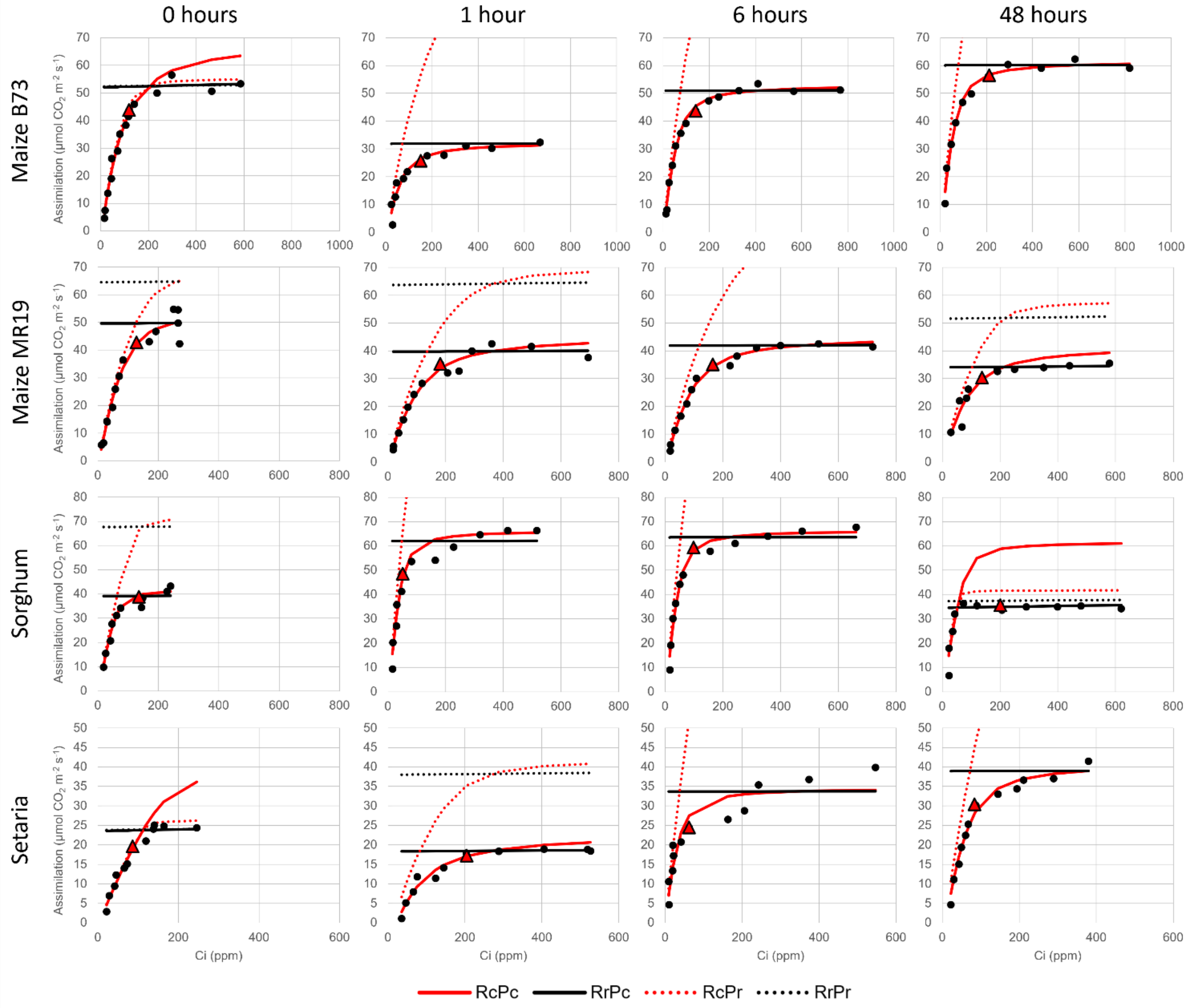


**Figure S2: Representative A/Ci curves.** Points are individual measurements. Triangles represent the measurement at 400ppm. RcPc: limited by RuBP carboxylation and PEP carboxylation. RrPc: limited by RuBP regeneration and PEP carboxylation. RcPr: limited by RuBP carboxylation and PEP regeneration. RrPr: limited by RuBP regeneration and PEP regeneration.

| | Maize B73 | | | | | | | | | | | | | | | | | | | |
|---|---|---|---|---|---|---|---|---|---|---|---|---|---|---|---|---|---|---|---|---|
| | 0 hours | | | | | 1 hour | | | | | 6 hours | | | | | 48 hours | | | | |
| | Plant 1 | Plant 2 | Plant 3 | Mean | Std Err | Plant 1 | Plant 2 | Plant 3 | Mean | Std Err | Plant 1 | Plant 2 | Plant 3 | Mean | Std Err | Plant 1 | Plant 2 | Plant 3 | Mean | Std Err |
| **Vcmax** | 67.2 | 87.8 | 70.2 | 75.1 | 6.42 | 65.7 | 34.8 | 40.6 | 47.0 | 9.49 | 57.9 | 58.2 | 67.2 | 61.1 | 3.08 | 68.8 | 58.8 | 57.9 | 61.8 | 3.49 |
| **J** | 799 | 331 | 494 | 541 | 137 | 714 | 68.3 | 719 | 500 | 216 | 764 | 743 | 912 | 806 | 53.4 | 963 | 658 | 588 | 736 | 115 |
| **Vpmax** | 276 | 98.7 | 114 | 163 | 56.8 | 212 | 20.5 | 166 | 133 | 57.7 | 231 | 244 | 243 | 239 | 4.44 | 280 | 273 | 245 | 266 | 10.8 |
| **Rd** | 10.0 | 10.0 | 10.0 | 10.0 | 0.00 | 2.75 | 5.75 | 6.02 | 4.84 | 1.05 | 2.65 | 2.93 | 7.40 | 4.33 | 1.54 | 4.84 | 6.04 | 0.10 | 3.66 | 1.81 |
| **gm** | 5.12 | 19.9 | 13.4 | 12.8 | 4.27 | 171 | 9.32 | 354 | 178 | 99.6 | 181 | 200 | 500 | 294 | 103 | 500 | 500 | 127 | 376 | 124 |
| **RMSE** | 367.6 | 67.7 | 58.3 | | | 32.2 | 4.08 | 64.6 | | | 53.8 | 26.6 | 15.5 | | | 56.5 | 59.3 | 45.7 | | |
| **Limit** | RcPc | RcPc | RcPc | | | RcPc | RcPc | RcPc | | | RcPc | RcPc | RcPc | | | RcPc | RcPc | RcPc | | |

| | Maize MR19 | | | | | | | | | | | | | | | | | | | | | |
|---|---|---|---|---|---|---|---|---|---|---|---|---|---|---|---|---|---|---|---|---|---|---|
| | 0 hours | | | | | | | 1 hour | | | | | 6 hours | | | | | 48 hours | | | | |
| | Plant 1 | Plant 2 | Plant 3 | Plant 4 | Plant 5 | Mean | Std Err | Plant 1 | Plant 2 | Plant 3 | Mean | Std Err | Plant 1 | Plant 2 | Plant 3 | Mean | Std Err | Plant 1 | Plant 2 | Plant 3 | Mean | Std Err |
| **Vcmax** | 50.6 | 48.1 | 87.7 | 67.5 | 80.5 | 66.9 | 7.85 | 61.5 | 51.9 | 41.6 | 51.7 | 5.74 | 52.8 | 57.8 | 63.6 | 58.0 | 3.11 | 56.6 | 33.3 | 44.9 | 44.9 | 6.73 |
| **J** | 327 | 260 | 326 | 388 | 344 | 329 | 20.7 | 676 | 366.3 | 337 | 460 | 108 | 480 | 531 | 693 | 568 | 64.3 | 168 | 792 | 293 | 418 | 191 |
| **Vpmax** | 153 | 69.0 | 119 | 108 | 84 | 107 | 14.5 | 475 | 127.8 | 115 | 239 | 118 | 150 | 164 | 185 | 166 | 10.2 | 343 | 524 | 134 | 334 | 113 |
| **Rd** | 9.86 | 10.0 | 9.95 | 8.78 | 10.0 | 9.72 | 0.24 | 10.00 | 2.24 | 2.97 | 5.07 | 2.47 | 4.12 | 0.05 | 0.05 | 1.41 | 1.36 | 10.00 | 0.05 | 0.05 | 3.37 | 3.32 |
| **gm** | 5.98 | 40.5 | 5.55 | 13.6 | 84.2 | 30.0 | 15.0 | 2.51 | 83.81 | 79.5 | 55.3 | 26.4 | 83.8 | 126.0 | 500.0 | 237 | 132 | 1.97 | 2.42 | 47.4 | 17.3 | 15.1 |
| **RMSE** | 42.1 | 235 | 261.9 | 136.5 | 216.6 | | | 180.9 | 50.76 | 19.6 | | | 19.4 | 12.3 | 43.3 | | | 44.5 | 27.7 | 78.4 | | |
| **Limit** | RcPc | RcPc | RcPc | RcPc | RcPc | | | RcPc | RcPc | RcPc | | | RcPc | RcPc | RcPc | | | RcPc | RcPc | RcPc | | |

| | Sorghum BTx623 | | | | | | | | | | | | | | | | | | | |
|---|---|---|---|---|---|---|---|---|---|---|---|---|---|---|---|---|---|---|---|---|
| | 0 hours | | | | | 1 hour | | | | | 6 hours | | | | | 48 hours | | | | |
| | Plant 1 | Plant 2 | Plant 3 | Mean | Std Err | Plant 1 | Plant 2 | Plant 3 | Mean | Std Err | Plant 1 | Plant 2 | Plant 3 | Mean | Std Err | Plant 1 | Plant 2 | Plant 3 | Mean | Std Err |
| **Vcmax** | 64.0 | 54.0 | 66.2 | 61.4 | 3.77 | 78.0 | 51.8 | 22.0 | 50.6 | 16.2 | 57.9 | 58.2 | 67.2 | 61.1 | 3.08 | 68.8 | 58.8 | 57.9 | 61.8 | 3.49 |
| **J** | 397 | 410 | 396 | 401 | 4.59 | 792 | 402 | 659 | 618 | 114 | 764 | 743 | 912 | 806 | 53.4 | 963 | 658 | 588 | 736 | 115 |
| **Vpmax** | 193 | 152 | 194 | 180 | 13.7 | 452 | 279 | 184 | 305 | 78.5 | 231 | 244 | 243 | 239 | 4.44 | 280 | 273 | 245 | 266 | 10.8 |
| **Rd** | 10.0 | 10.0 | 10.0 | 10.0 | 0.00 | 10.0 | 7.41 | 1.05 | 6.15 | 2.66 | 2.65 | 2.93 | 7.40 | 4.33 | 1.54 | 4.84 | 6.04 | 0.10 | 3.66 | 1.81 |
| **gm** | 9.56 | 13.9 | 9.19 | 10.9 | 1.52 | 500 | 500 | 500 | 500 | 0.00 | 181 | 200 | 500 | 294 | 103 | 500 | 500 | 127 | 376 | 124 |
| **RMSE** | 207 | 55.0 | 236 | | | 220 | 48.0 | 13.3 | | | 53.8 | 26.6 | 15.5 | | | 56.5 | 59.3 | 45.7 | | |
| **Limit** | RcPc | RrPc | RcPc | | | RcPc | RcPc | RcPc | | | RcPc | RcPc | RcPc | | | RrPc | RrPc | RrPc | | |

| | Setaria A10 | | | | | | | | | | | | | | | | | | | |
|---|---|---|---|---|---|---|---|---|---|---|---|---|---|---|---|---|---|---|---|---|
| | 0 hours | | | | | 1 hour | | | | | 6 hours | | | | | 48 hours | | | | |
| | Plant 1 | Plant 2 | Plant 3 | Mean | Std Err | Plant 1 | Plant 2 | Plant 3 | Mean | Std Err | Plant 1 | Plant 2 | Plant 3 | Mean | Std Err | Plant 1 | Plant 2 | Plant 3 | Mean | Std Err |
| **Vcmax** | 51.2 | 48.4 | 79.3 | 59.6 | 9.88 | 48.5 | 35.8 | 44.7 | 43.0 | 3.76 | 65.2 | 35.8 | 30.3 | 43.8 | 10.9 | 53.2 | 64.6 | 49.2 | 55.6 | 4.60 |
| **J** | 157 | 160 | 171 | 163 | 4.17 | 792 | 263 | 792 | 616 | 176 | 664 | 792 | 445 | 634 | 101 | 667 | 456 | 883 | 669 | 123 |
| **Vpmax** | 65.7 | 98.2 | 91.0 | 84.9 | 9.86 | 272 | 104 | 325 | 234 | 66.6 | 443 | 280 | 158 | 294 | 82.4 | 227 | 207 | 271 | 235 | 19.0 |
| **Rd** | 0.05 | 5.38 | 10.0 | 5.14 | 2.87 | 7.61 | 10.0 | 6.82 | 8.14 | 0.96 | 10.0 | 0.05 | 0.05 | 3.37 | 3.32 | 10.0 | 10.0 | 9.27 | 9.76 | 0.24 |
| **gm** | 229 | 3.33 | 2.84 | 78.3 | 75.2 | 500 | 254 | 260 | 338 | 81.0 | 500 | 500 | 134 | 378 | 122 | 52.9 | 130 | 135 | 106 | 26.5 |
| **RMSE** | 155 | 20.3 | 89.5 | | | 62.5 | 15.7 | 155 | | | 83.4 | 177 | 148 | | | 35.1 | 233 | 141 | | |
| **Limit** | RcPc | RcPc | RcPr | | | RcPc | RcPc | RcPc | | | RcPc | RcPc | RcPc | | | RcPc | RcPc | RcPc | | |

**Table S1: A/Ci curve fit values.** RMSE is root mean square error from the data points. Curves that do not follow the RcPc limitation are noted with red text. RcPc: limited by RuBP carboxylation and PEP carboxylation. RrPc: limited by RuBP regeneration and PEP carboxylation. RcPr: limited by RuBP carboxylation and PEP regeneration. RrPr: limited by RuBP regeneration and PEP regeneration.

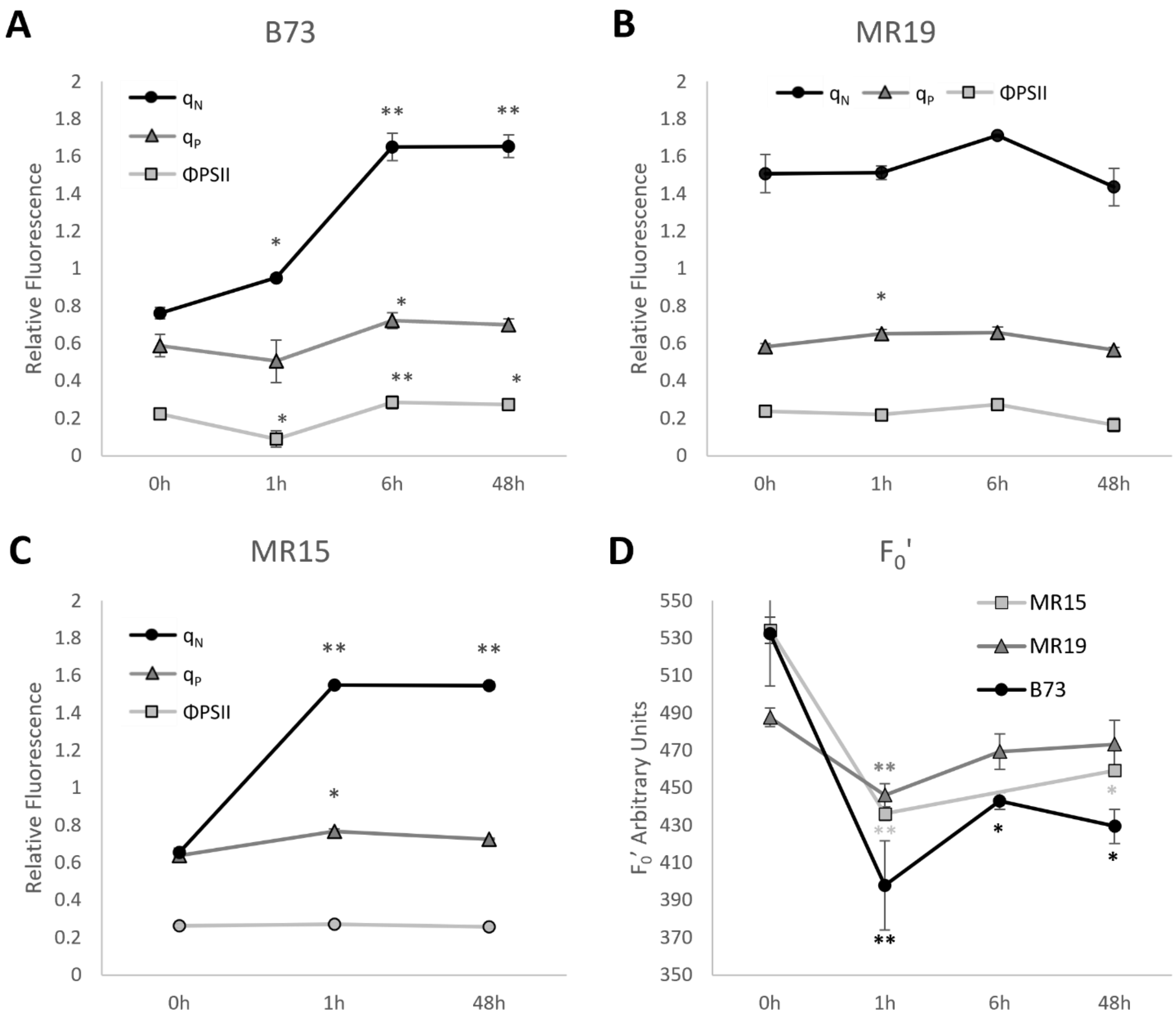


**Figure S3**: Chlorophyll fluorescence data for select maize cultivars. **A-C.** Photon partitioning for maize B73 (A), MR19 (B) and MR15 (C). $q_N$ nonphotochemical quenching parameter, $q_P$ photochemical quenching parameter, ΦPSII quantum yield of photosystem II. **D.** Light-adapted basal fluorescence parameter ($F_0$') of all three cultivars. * $p<0.05$, ** $p<0.01$ vs. 0-hour control. Error bars are standard error of at least 3 biological replicates.

MR15 (Figure S3C) displays the typical result of heat exposure, with $q_N$ rapidly increasing and remaining elevated throughout the duration of treatment. $q_P$ and ΦPSII remain largely unchanged. In contrast, B73 has delayed induction of $q_N$, which remains fairly low at 1 hour (Figure S3A). This change is correlated with a transient decrease in ΦPSII. $q_N$ is fully induced by 6 hours and remains elevated at 48 hours, while ΦPSII is slightly higher than the control value at both time points. Interestingly, MR19 displays elevated $q_N$ even at the 0h condition of 25$^o$C (Figure S3B). No changes to $q_N$, $q_P$ or ΦPSII are seen in MR19 during the heat treatment.

Damage caused by high temperature generally results in an increase in $F_0$', while acceptor-side limitations such as $CO_2$ assimilation result in decreased $F_0$' (Sharkey et al., 2001). All three cultivars show significantly reduced $F_0$' at 1 hour of heat, indicating that $CO_2$ assimilation is limiting at that time point (Figure S3D).

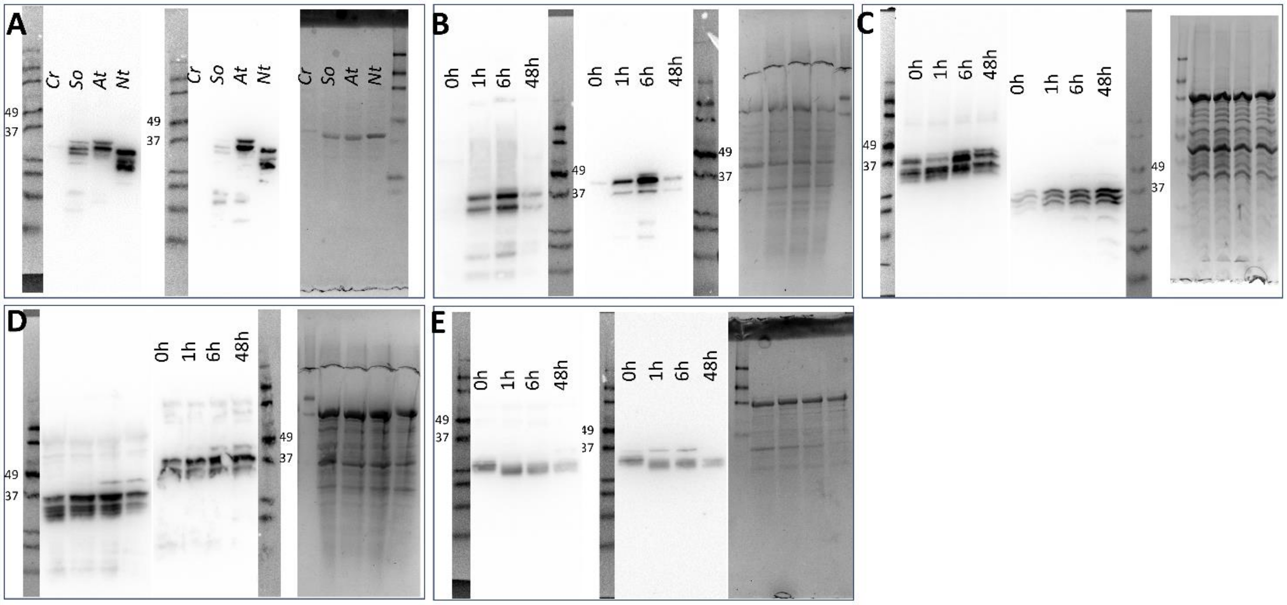


**Figure S4**: complete Western blots. **A**. controls. **B.** Maize B73. **C.** Maize MR19. **D.** Sorghum. **E.** Setaria. Each panel, left to right: Agrisera antibody, Huabio antibody, Coomassie loading control (panel A, purified protein; panels B-E, total protein before purification). Positions of the 49kDa and 37kDa markers are indicated.

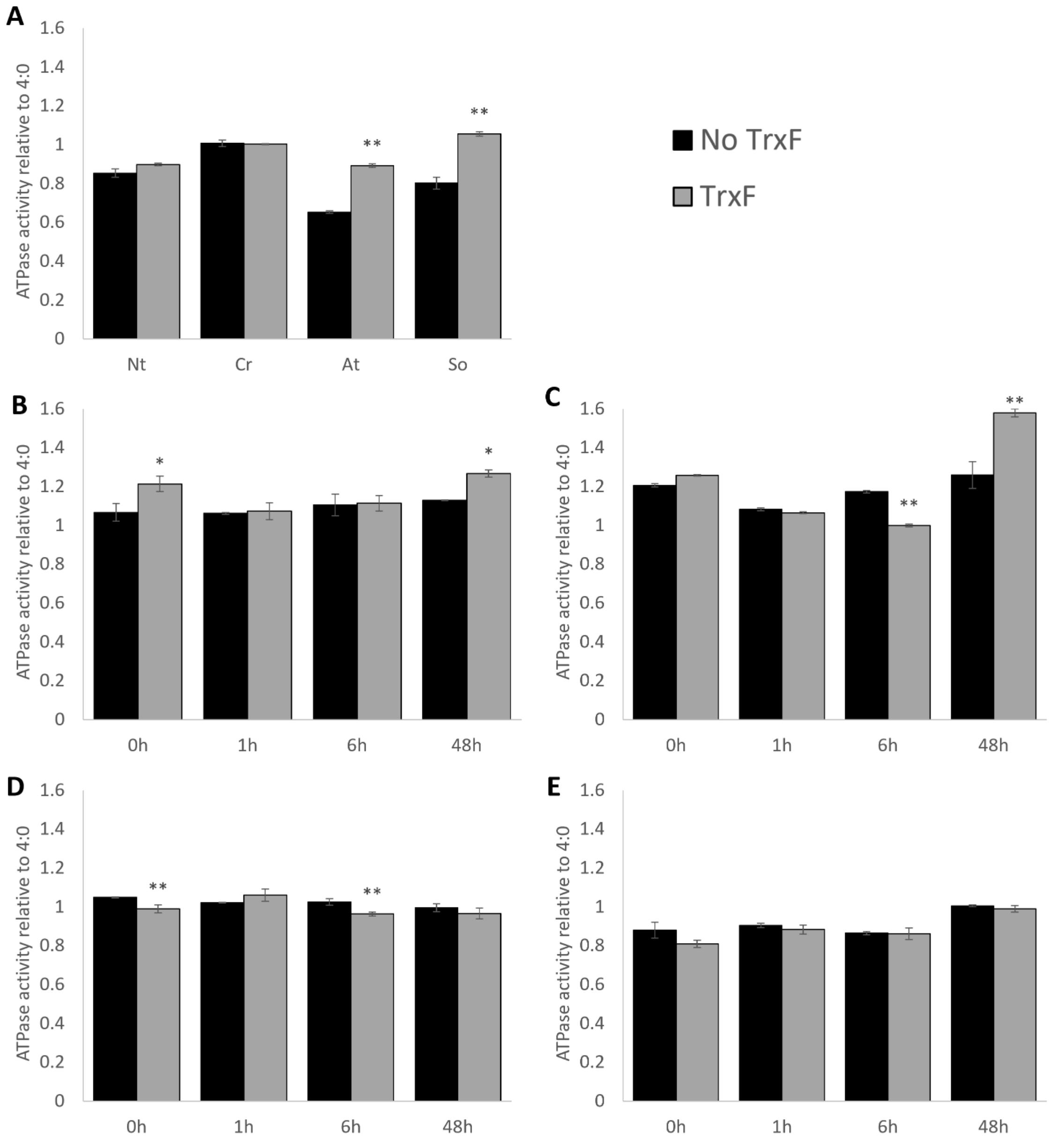


**Fig S5**: response of ATPase activity to TrxF relative to 4:0 ratio. ATP:ADP ratio was 3:1 and 10mM $Mg^{2+}$. **A**. Controls **B**. maize B73 **C**. maize MR19 **D**. sorghum **E**. setaria. * p<0.05, ** p<0.01 vs. no TrxF, error bars are standard deviation of n=5

We investigated the influence of redox control of RCA activity in the different C4 plants. RCA contains a disulfide bond present in the CTD of the α isoform, which is reduced by thioredoxin F (TrxF). We therefore tested RCA ATPase activity in the presence and absence of reduced TrxF. We used RCA proteins from Arabidopsis, spinach, tobacco and Chlamydomonas to reproduce published results: stimulation of ATPase activity in Arabidopsis and spinach by TrxF, and the lack of stimulation in both tobacco and Chlamydomonas, both of which lack the α isoform. TrxF-dependent stimulation of ATPase activity was weakly observed in the maize cultivars but does not appear to be relevant in sorghum or setaria.

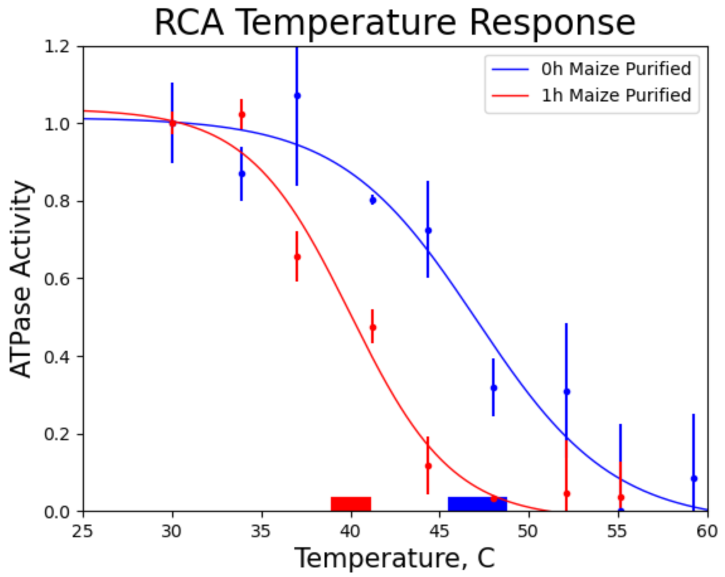


**Figure S6**: a representative temperature response curve for maize B73 0h and 1h samples. $T_{50}$ values are represented by bars at the bottom of the graph. Error bars are standard deviation, n=3.

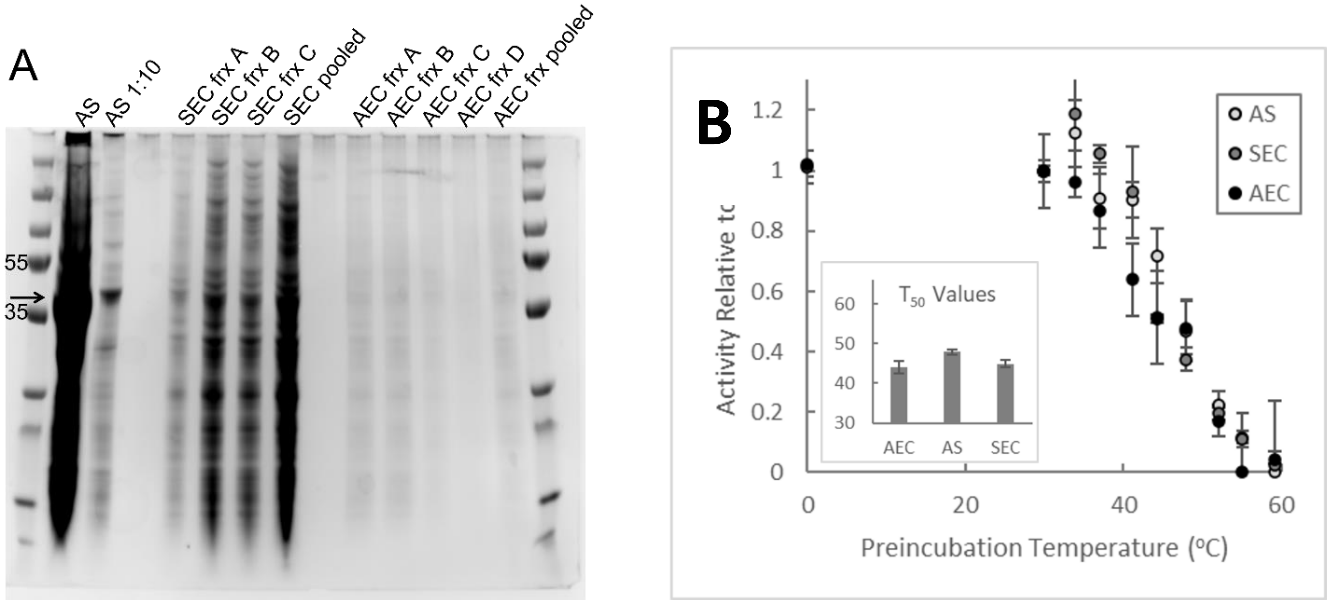


**Figure S7**: Comparison of RCA purification methods. **A.** total protein stain for maize tissue purified with ammonium sulfate precipitation only (AS); ammonium sulfate precipitation and size exchange chromatography (SEC); or ammonium sulfate precipitation, size exchange chromatography and anion exchange chromatography (AEC), multiple fractions (frx) shown for SEC and AEC samples. Expected RCA size indicated by arrow. **B.** Thermostability assay on AS, SEC and AEC samples, with calculated $T_{50}$ values in inset. Mean and standard deviation of 3 technical replicates. Differences in $T_{50}$ were not significant.
